# The MRL/MpJ Mouse Strain Is Not Protected From Muscle Atrophy And Weakness After Rotator Cuff Tear

**DOI:** 10.1101/702167

**Authors:** Jeffrey R Talarek, Alex N Piacentini, Alexis C Konja, Susumu Wada, Jacob B Swanson, Samuel C Nussenzweig, Joshua S Dines, Scott A Rodeo, Christopher L Mendias

**Affiliations:** Hospital for Special Surgery, New York, NY; Departments of Orthopaedic Surgery, Weill Cornell Medical College, New York, NY; Departments of Physiology and Biophysics, Weill Cornell Medical College, New York, NY

**Author notes:** Corresponding Author Christopher Mendias, PhD, Hospital for Special Surgery 535 E 70th St New York, NY 10021 USA, office.

**Keywords:** muscle atrophy, myosteatosis, tenotomy, metabolomics, lipidomics

## Abstract

Chronic rotator cuff tears are a common source of shoulder pain and disability. Patients with rotator cuff tears often have substantial weakness, fibrosis, and fat accumulation which limit successful surgical repair and postoperative rehabilitation. The Murphy Roths Large (MRL) strain of mice have demonstrated superior healing and protection against pathological changes in several disease and injury conditions. We tested the hypothesis that, compared to the commonly used C57Bl/6 (B6) strain, MRL mice would have less muscle fiber atrophy and fat accumulation, and be protected against the loss in force production that occurs after cuff tear. Adult male mice B6 and MRL mice were subjected to a rotator cuff tear, and changes in muscle fiber contractility and histology were measured. RNA sequencing, and shotgun metabolomics and lipidomics were also performed. Muscles were harvested one month after tear. B6 and MRL mice had a 40% reduction in relative muscle force production after rotator cuff tear. RNA sequencing identified an increase in fibrosis-associated genes and a reduction in mitochondrial metabolism genes. Markers of glycolytic metabolism increased in B6 mice, while MRL mice appeared to increase amino acid metabolism after tear. There was an accumulation of lipid after injury, although there was a divergent response between B6 and MRL mice in the types of lipid species that accrued. There were strain-specific differences between the transcriptome, metabolome, and lipidome of B6 and MRL mice, but these differences did not protect MRL mice from weakness and pathological changes after rotator cuff tear.

## Introduction

Rotator cuff tears are among the most frequent upper extremity injuries, with over a quarter of a million surgical repairs performed in the US on an annual basis ^1^. While patients are generally satisfied with pain relief after rotator cuff repair, many remain unsatisfied with persistent muscle weakness ^2^. The strength deficits that patients experience are due to the pathological changes that occur in torn rotator cuff muscles, including fibrosis, muscle fiber atrophy, and fat accumulation in and around muscle fibers ^3–5^. These combined pathological changes, often referred to as fatty degeneration or myosteatosis, are correlated with reduced muscle force production and are a limiting factor in achieving successful surgical repair and restoring strength and function through post-operative rehabilitation ^3,6,7^. Gaining a greater understanding of the biological mechanisms that lead to fatty degeneration could identify novel therapies to improve clinical outcomes for patients with chronic rotator cuff tears.

The Murphy Roths Large (MRL/MpJ, or MRL) line of mice have received attention as a "super healing" strain with enhanced tissue regeneration compared to many standard mouse strains ^8^. MRL mice have an absence of articular cartilage degeneration and improved regeneration and after joint injury ^9,10^, improved mechanical properties and healing after tendon injury, and reduced muscle fibrosis in a γ-sarcoglycan-deficient model of muscular dystrophy ^11–13^. However, the MRL strain does not display enhanced tissue healing for every organ system and type of injury, with no differences noted between MRL mice and controls in dorsal skin healing following punch injury, nor in cardiac healing following infarct ^14,15^.

Given the context-dependent improvement in tissue healing in MRL mice, we sought to compare the pathological changes that occur in MRL mice following a rotator cuff tear. Based on the improvement in healing and reduced pathological changes in other musculoskeletal injury and disease models, we formulated the hypothesis that MRL mice would display less muscle fiber atrophy and fat accumulation, and experience less of a loss in strength, than C57Bl/6 (B6) mice. To test this hypothesis we performed a full-thickness supraspinatus and infraspinatus tenectomy and denervation in adult male MRL and B6 mice, and measured muscle fiber contractility and histology. We also performed RNA sequencing (RNAseq) as well as mass spectrometry-based muscle metabolomics and lipidomics to further explore biological differences between B6 and MRL mice after a rotator cuff tear.

## Methods

### Animals

This study was approved by the Hospital for Special Surgery/Weill Cornell/Memorial Sloan Kettering IACUC (protocol 2017-0035). All experiments were performed in accordance with the approved protocol. Three-month old male C57Bl/6J (B6, strain 000664, Jackson Labs, Bar Harbor, ME) and MRL/MpJ (MRL, strain 000486, Jackson Labs) were used. Mice were housed under specific pathogen free conditions and allowed *ad libidum* access to food and water, but were fasted for four hours prior to harvest to minimize any impacts of acute food consumption on metabolomic measurements. A total of N=20 mice were used in the study, with N=4 control B6 mice, N=4 control MRL mice, N=6 surgical B6 mice, and N=6 surgical MRL mice.

### Surgical Model

A bilateral, full-thickness supraspinatus and infraspinatus tenectomy and denervation were performed as modified from previous studies ^16,17^. Mice were anesthetized with 2% isoflurane, and a deltoid-splitting transacromial approach was used to expose the supraspinatus and infraspinatus tendons which were then finely detached from their footprint on the humerus. Tendons were then transected just distal to the myotendinous junction, allowing for removal of the tendon without directly causing damage to the muscle during the procedure, in order to prevent spontaneous reattachment of the tendon to the surrounding fascia ^18,19^. The suprascapular nerve was identified, and an approximately 3mm segment was removed to denervate the muscles. Denervation was performed as biopsies from patients with rotator cuff tears demonstrate signs of muscle fiber denervation ^20^. The incision was closed with interrupted sutures using 6-0 Prolene (Ethicon, Somerville, NJ) and animals were allowed to recover in their cages. Mice were treated with buprenorphine twice post-operatively (0.05mg/kg, Reckitt, Parsippany, NJ).

After a period of 30 days, mice were anesthetized with 2% isoflurane, and supraspinatus and infraspinatus muscles were harvested and weighed. The left supraspinatus was prepared for muscle fiber contractility measurements. The right supraspinatus was mounted on a cork disk with tragacanth gum, snap frozen in isopentane and stored at −80°C for histology. The left infraspinatus was finely minced and equal portions were divided and snap frozen in liquid nitrogen and stored at −80°C to be used for metabolomics or lipidomics studies. The right infraspinatus was snap frozen and stored at −80°C for RNA isolation. Soleus muscles were also harvested to verify fiber type antibody staining. Animals were humanely euthanized by cervical dislocation.

### Muscle Fiber Contractility

Permeabilized muscle fiber contractility was performed as modified from previous publications ^16,21^. Small bundles of muscle fibers were dissected free of the muscle, chemically permeabilized, and stored at −80°C. Individual fibers were placed in a chamber containing relaxing solution and secured at one end to a servomotor (Aurora Scientific, Aurora, ON) and the other end to a force transducer (Aurora Scientific) with two ties of 10-0 monofilament nylon suture. Using a video sarcomere length measurement system (Aurora Scientific), fiber length was adjusted to obtain a sarcomere length of 2.5µm. The average fiber CSA was calculated assuming the fiber had an elliptical cross-section, with diameters obtained at five positions along the length of the fiber from top and side views. Maximum isometric force (F_o_) was measured by immersing the fiber in a high calcium activation solution, and specific maximum isometric force (sF_o_) was determined by dividing F_o_ by fiber CSA. Eight to ten fibers were tested from each supraspinatus muscle. Due to the nature of the permeabilization process, fibers swell in proportion to their CSA ^22^, and therefore CSA values of permeabilized fibers are greater than measurements from histology.

### Histology

Histology was performed as described ^23,24^. Sections were blocked with a MOM Blocking Kit (Vector Labs, Burlingame, CA) and incubated with primary antibodies from Developmental Studies Hybridoma Bank (Iowa City, IA) against type I myosin heavy chain (MHC) (BA-D5), type IIA MHC (SC-71), and type IIB MHC (BF-F3). Type IIX MHC was identified by the lack of signal within the outline of a fiber. Secondary antibodies conjugated to either AlexaFluor 350, 488, or 555 fluorescent probes (Thermo Fisher Scientific, Waltham, MA) were used to detect primary antibodies. Glycosaminoglycans (GAGs) in the extracellular matrix were detected using wheat germ agglutinin (WGA) conjugated to AlexaFluor 647. Digital images of slides were captured using an EVOS FL microscope (Thermo Scientific). Cross-sectional areas (CSA) of muscle fibers were measured with ImageJ (NIH, Bethesda, MD).

### Gene Expression and RNA Sequencing

RNA was isolated from muscles, and gene expression and RNA sequencing (RNAseq) was performed as described ^16^. Samples were homogenized in QIAzol (Qiagen, Valencia, CA) and RNA was isolated using 5PRIME Phase Lock Gel tubes (Quantabio, Beverly, MA), an RNeasy Mini kit (Qiagen), and DNase I (Qiagen). RNA concentration and quality were assessed using a NanoDrop One (Thermo Fisher Scientific) and a BioAnalyzer 2100 (Agilent, Santa Clara, CA). All samples had RIN values greater than 8.1.

For RNAseq, a total of 250ng of RNA was delivered to the Weill Cornell Epigenomics Core for processing and sequencing. RNA sample concentrations were normalized and cDNA pools were created for each sample, and then subsequently tagged with a barcoded oligo adapter to allow for sample-specific resolution. Sequencing was carried out using an Illumina HiSeq 2500 platform (Illumina, San Diego, CA) with 50bp single end reads. Data was quality checked using FastQC, and aligned to the reference genome (mm10, UCSC, Santa Cruz, CA). Differential expression was calculated using edgeR ^25^. Complete RNAseq data has been deposited to NIH GEO (ascension number GSE130447).

To validate select differentially expressed genes from RNAseq, quanitative PCR (qPCR) was performed. RNA was reversed transcribed into cDNA using iScript Reverse Transcription Supermix (Bio-Rad). Quantitative PCR (qPCR) was performed with cDNA in a CFX96 real-time thermal cycler (Bio-Rad) using SsoFast SYBR Green Supermix (Bio-Rad). Target gene expression was normalized to the housekeeping gene Casein Kinase 2α1 (*Csnk2a1*) using the 2^−ΔCt^ method.

### Biological Pathway Analysis

Expression data from RNAseq measurements was imported into Ingenuity Pathway Analysis (IPA) software (Qiagen) to assist in predicting cellular and molecular pathways that were altered between strains, and before and after rotator cuff tears.

### Metabolomics and Lipidomics

Mass spectrometry-based metabolomics and lipidomics was performed by the University of Michigan Metabolomics Core as described ^16,26^. Summary data is provided in the manuscript, with complete data available in Supplemental Tables S1 and S2. For metabolomics, metabolites were extracted from frozen muscle, derivatized, and analyzed with gas chromatography-MS ^27^. Quantification of metabolites was performed using Masshunter Quantitative Analysis software (Agilent Technologies, Santa Clara, CA, USA). Two samples from the B6 tear group failed quality control and were excluded from analysis.

For lipidomics, lipids were extracted from samples and subjected to liquid chromatrography-mass spectrometry (LC-MS). MS peaks were matched in-silico with LipidBlast ^28^. Quantification was performed by Multiquant software (AB-SCIEX, Framingham, MA). One WT sample failed quality control and was excluded from analysis. MetaboAnalyst 4.0 ^29^ was used to perform principal component (PC) analyses.

### Statistics

Values are presented as mean±standard deviation (SD). Differences between groups were determined using a two-way ANOVA (α=0.05) followed by Fisher’s LSD post-hoc sorting. Differences in fiber type distribution were assessed with a chi-squared test. For RNAseq, metabolomics, and lipidomics data, a false discovery rate correction was used to adjust p-values for multiple observations, and are reported as q-values. With the exception of analyses performed in edgeR, statistical calculations were performed in Prism software (version 8, GraphPad, San Diego, CA).

## Results

Following rotator cuff tear, the body weight normalized supraspinatus muscle mass of B6 mice was reduced by nearly 50%, while MRL mice displayed a 42% reduction in mass (Table 1). Infraspinatus muscles demonstrated a similar trend (Table 1). Neither strain lost body mass after rotator cuff tear, although the MRL mice were 55% larger than B6 mice at baseline (Table 1).

**Table 1.** Body and muscle mass data. Values are mean±SD. Differences between groups assessed using a two-way ANOVA. Posthoc sorting: a, different (p<0.05) from B6 control; b, different (p<0.05) from B6 tear; c, different (p<0.05) from MRL control. N≥4 mice per group.

|  | **B6** |  | **MRL** |  |
| --- | --- | --- | --- | --- |
|  | *Control* | *Tear* | *Control* | *Tear* |
| Body mass (g) | 30.4±1.8 | 27.1±1.2 | 47.2±1.6<sup>a,b</sup> | 47.1±2.5<sup>a,b</sup> |
| Supraspinatus mass (mg) | 37.1±1.5 | 17.1±2.2<sup>a</sup> | 54.3±3.5<sup>a,b</sup> | 31.4±2.6<sup>a,b,c</sup> |
| Supraspinatus/body mass (mg/g) | 1.22±0.08 | 0.62±0.07<sup>a</sup> | 1.15±0.04<sup>b</sup> | 0.67±0.05<sup>a,c</sup> |
| Infraspinatus mass (mg) | 33.0±4.2 | 16.8±1.8<sup>a</sup> | 50.3±6.2<sup>a,b</sup> | 30.9±2.5<sup>b,c</sup> |
| Infraspinatus/body mass (mg/g) | 1.08±0.10 | 0.61±0.07<sup>a</sup> | 1.06±0.10<sup>b</sup> | 0.66±0.07<sup>a,c</sup> |

To assess functional changes in rotator cuff muscles after injury, we measured permeabilized muscle fiber contractility. No changes in permeabilized fiber CSA were noted (Figure 1A). F_o_ of control MRL mice was 44% larger than control B6 mice, and after tear F_o_ was reduced in B6 mice by 46% and by 50% in MRL mice (Figure 1B). sF_o_ was 52% higher in control MRL mice than B6 mice, and both groups were reduced by 40% after tear (Figure 1C).

**Figure 1.**
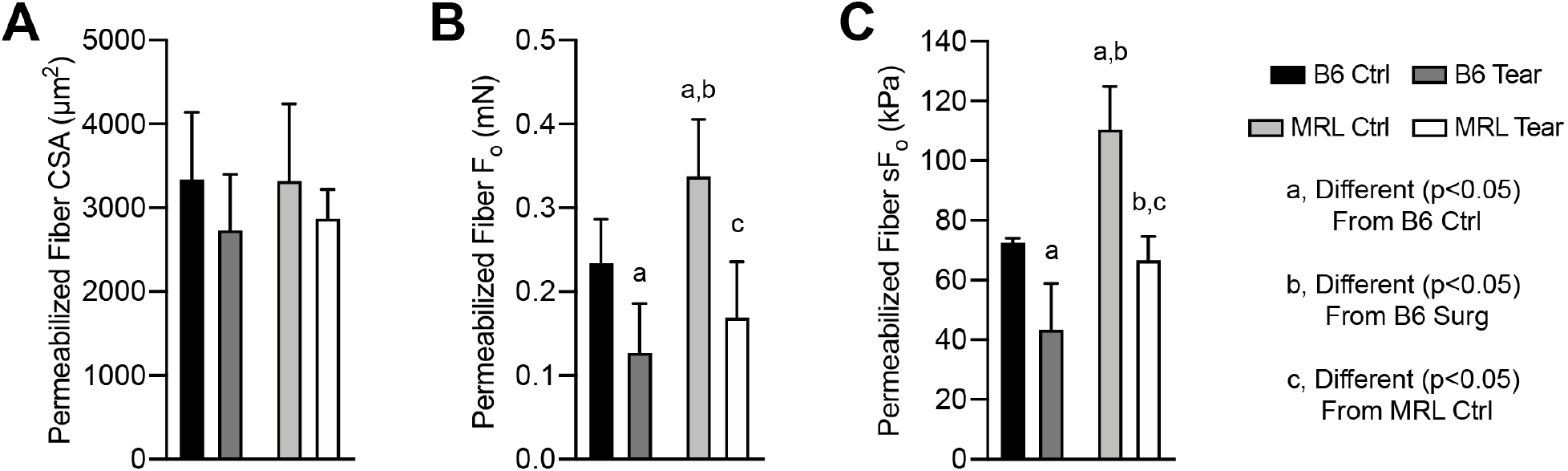
Muscle Fiber Contractility. (A) Permeabilized muscle fiber cross-sectional area (CSA), (B) maximum isometric force (F_o_), and (C) specific maximum isometric force (sF_o_) of supraspinatus muscles. Black, B6 control (Ctrl); dark gray, B6 tear; light gray, MRL ctrl; white, MRL tear. Differences between groups assessed using a two-way ANOVA. Posthoc sorting: a, different (p<0.05) from B6 control; b, different (P<0.05) from B6 tear; c, different (p<0.05) from MRL control. N≥4 muscles per group.

We next analyzed changes in histology. We detected type IIA, IIB, IIX, IIA/IIX, and IIB/IIX fibers in rotator cuff muscles, but did not observe type I fibers (Figure 2A). Soleus muscles of B6 mice, which are known to contain type I fibers, were used as a positive control (Figure 2A). There was a significant difference in the distribution of fiber types between B6 control and B6 tear muscles, MRL control and MRL tear muscles, and B6 tear and MRL tear muscles, but no difference in distribution between B6 control and MRL control muscles (Figure 2B). The most abundant fiber type in all conditions were type IIB fibers. However, for B6 mice the percentage of IIB fibers went from 58% to 45%, and IIX fibers increased in proportion from 27% to 35% (Figure 2B). In MRL mice, IIB fibers dropped from 50% to 36% after tear, and the largest increase in fiber type was type IIA, going from 8% to 17% after tear (Figure 2B). Both strains lost fibers after injury, with B6 mice losing 36% of the muscle fibers per supraspinatus, while MRL mice lost 50% of the fibers per supraspinatus (Figure 2C). There were also differences in the size of fibers. The type IIB and IIX fibers of control MRL mice were 41% and 74% larger than B6, respectively (Figure 2D). After injury, the size of type IIB fibers was reduced by 38% in B6 mice and by 45% in MRL mice (Figure 2D).

**Figure 2.**
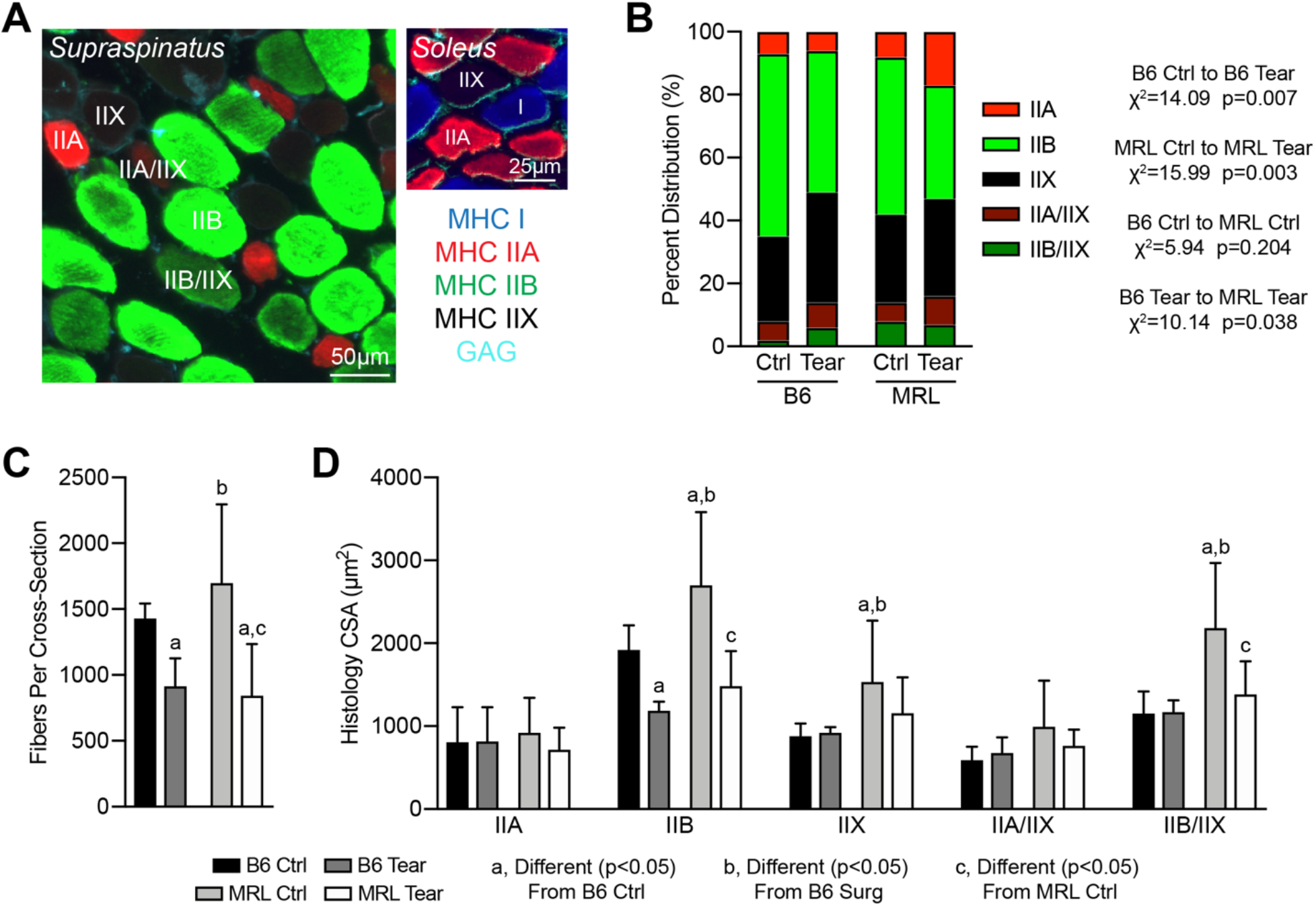
Fiber Type Histology. (A) Representative fiber type immunofluorescence labeling of supraspinatus muscles. A soleus muscle is shown as a control. Blue, myosin heavy chain (MHC) I; red, MHC IIA; green, MHC IIB; black (lack of signal), MHC IIX; glycosaminoglycans (GAG), teal. Scale bar for supraspinatus is 50µm and for soleus is 25µm. (B) Percent distribution by fiber type and chi-squared analysis. (C) Number of fibers per histology cross-section, and (D) cross-sectional area (CSA) of fibers by fiber type. Black, B6 control (Ctrl); dark gray, B6 tear; light gray, MRL ctrl; white, MRL tear. Differences between groups assessed using (B) a chi-squared test or (C-D) a two-way ANOVA. Posthoc sorting (C-D): a, different (p<0.05) from B6 control; b, different (p<0.05) from B6 tear; c, different (p<0.05) from MRL control. N≥4 muscles per group.

To identify global changes in the transcriptome of muscles, we performed RNAseq. We used IPA for pathway enrichment analysis and to identify genes that were important in the response to rotator cuff tear within and between strains (Table 2). To allow for detailed exploratory analysis, RNAseq is presented in two Figures. Figure 3 presents the differences between B6 tear to B6 control, MRL tear to MRL control groups, and individual genes are graphed based on known roles in fatty degeneration, and those identified from IPA analysis. Figure 4 shows the differences between B6 control to MRL control, B6 tear to MRL tear, and the top 30 genes that were upregulated and downregulated across strains and conditions.

**Table 2.**
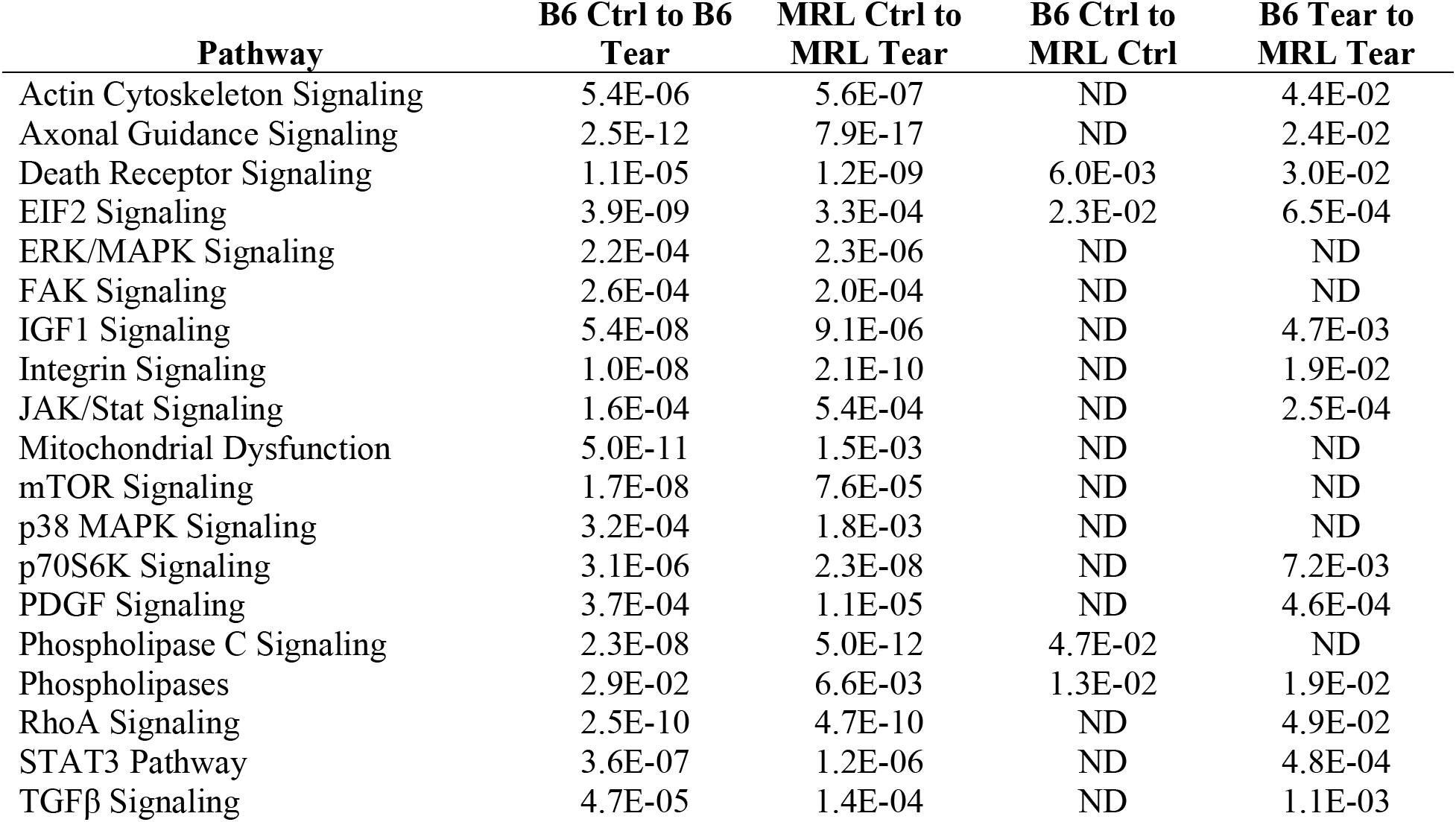
Gene enrichment analysis. The q-values of selected differentially regulated pathways that were identified using Ingenuity Pathway Analysis. ND, not significantly different.

| **Pathway** | **B6 Ctrl to B6 Tear** | **MRL Ctrl to MRL Tear** | **B6 Ctrl to MRL Ctrl** | **B6 Tear to MRL Tear** |
| --- | --- | --- | --- | --- |
| Actin Cytoskeleton Signaling | 5.4E-06 | 5.6E-07 | ND | 4.4E-02 |
| Axonal Guidance Signaling | 2.5E-12 | 7.9E-17 | ND | 2.4E-02 |
| Death Receptor Signaling | 1.1E-05 | 1.2E-09 | 6.0E-03 | 3.0E-02 |
| EIF2 Signaling | 3.9E-09 | 3.3E-04 | 2.3E-02 | 6.5E-04 |
| ERK/MAPK Signaling | 2.2E-04 | 2.3E-06 | ND | ND |
| FAK Signaling | 2.6E-04 | 2.0E-04 | ND | ND |
| IGF1 Signaling | 5.4E-08 | 9.1E-06 | ND | 4.7E-03 |
| Integrin Signaling | 1.0E-08 | 2.1E-10 | ND | 1.9E-02 |
| JAK/Stat Signaling | 1.6E-04 | 5.4E-04 | ND | 2.5E-04 |
| Mitochondrial Dysfunction | 5.0E-11 | 1.5E-03 | ND | ND |
| mTOR Signaling | 1.7E-08 | 7.6E-05 | ND | ND |
| p38 MAPK Signaling | 3.2E-04 | 1.8E-03 | ND | ND |
| p70S6K Signaling | 3.1E-06 | 2.3E-08 | ND | 7.2E-03 |
| PDGF Signaling | 3.7E-04 | 1.1E-05 | ND | 4.6E-04 |
| Phospholipase C Signaling | 2.3E-08 | 5.0E-12 | 4.7E-02 | ND |
| Phospholipases | 2.9E-02 | 6.6E-03 | 1.3E-02 | 1.9E-02 |
| RhoA Signaling | 2.5E-10 | 4.7E-10 | ND | 4.9E-02 |
| STAT3 Pathway | 3.6E-07 | 1.2E-06 | ND | 4.8E-04 |
| TGFβ Signaling | 4.7E-05 | 1.4E-04 | ND | 1.1E-03 |

**Figure 3.**
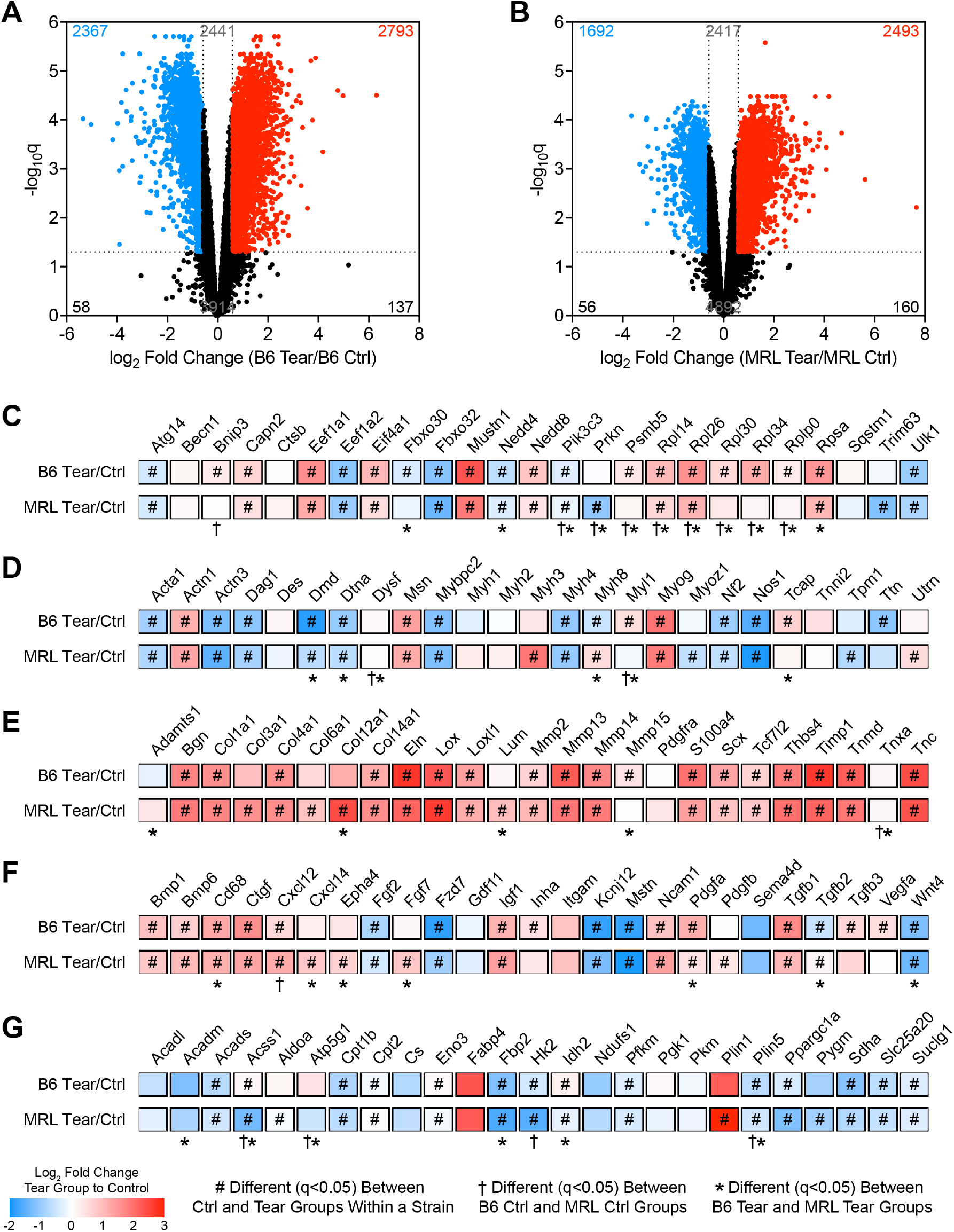
RNA Sequencing Comparing B6 Tear to MRL Control and MRL Tear to MRL Control. Volcano plot of transcripts that were differentially regulated between (A) B6 tear and control (ctrl) groups, and (B) MRL tear and ctrl groups. Heatmaps demonstrating the log_2_-fold change in tears to controls for both B6 and MRL mice for transcripts related to (C) hypertrophy and atrophy, (D) muscle fiber contractility and cytoskeletal organization, (E) extracellular matrix, (F) growth factors, cytokines, and receptors, and (G) metabolism. #, different (q<0.05) between ctrl and tear groups within a strain; †, different (q<0.05) between B6 ctrl and MRL ctrl groups; *, different (q<0.05) between B6 tear and MRL tear groups. N=4 infraspinatus muscles per group.

**Figure 4.**
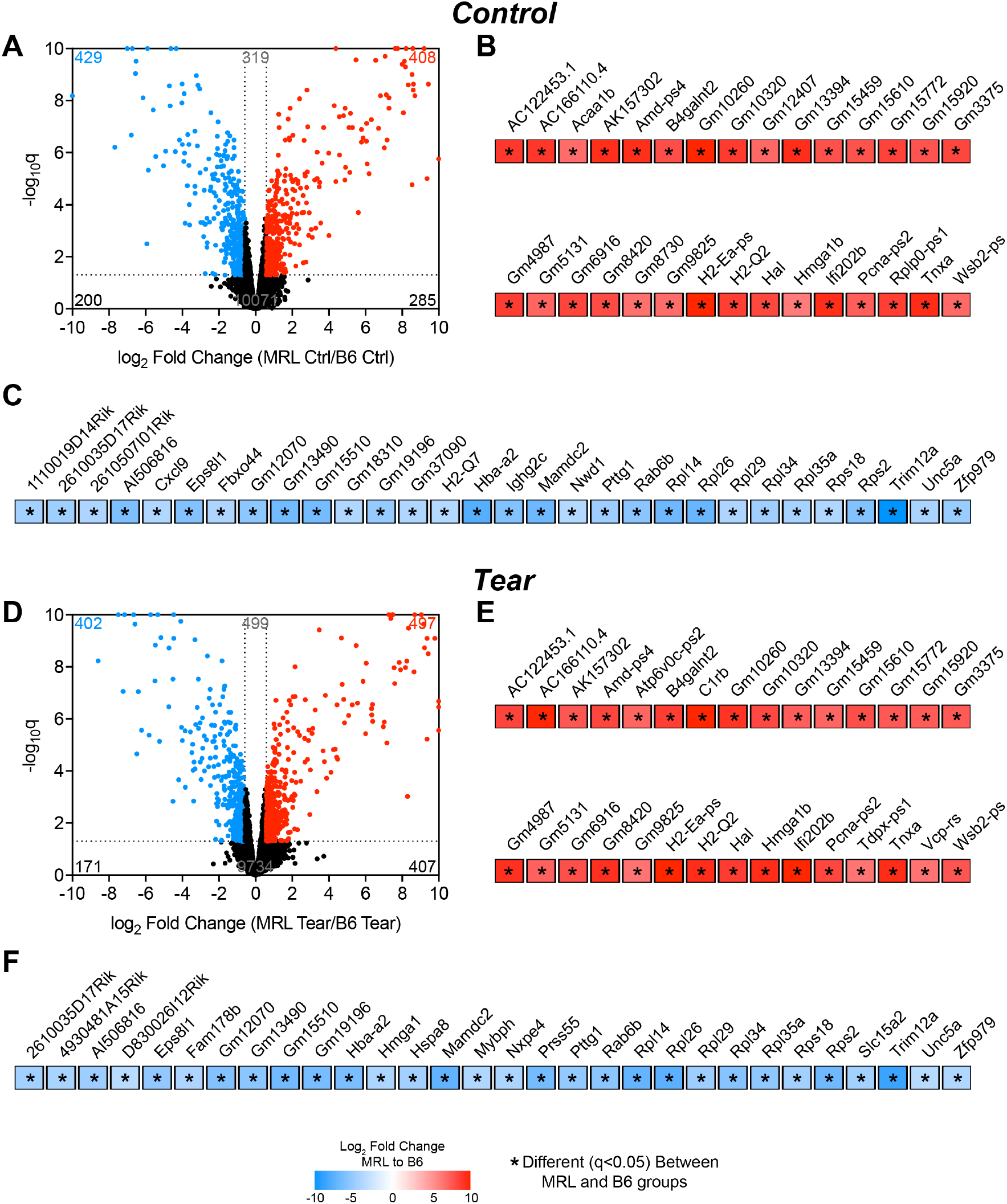
RNA Sequencing Comparing B6 Control to MRL Control and B6 Tear to MRL Tear. (A) Volcano plot of differentially regulated transcripts, and the top 30 (B) upregulated and (C) downregulated transcripts of B6 control and MRL control groups. (D) Volcano plot of differentially regulated transcripts, and the top 30 (E) upregulated and (F) downregulated transcripts of B6 tear and MRL tear groups. Data is expressed as log_2_-fold change of MRL groups normalized to B6 groups. *, different (q<0.05) between B6 tear and MRL tear groups. N=4 infraspinatus samples per group.

There were 2793 transcripts significantly upregulated by 50% in B6 tear muscles compared to B6 controls, and 2367 significantly downregulated by 50% (Figure 3A). In MRL mice, there were 2493 transcripts significantly upregulated by 50% and 1692 transcripts downregulated by 50% in tear groups compared to controls (Figure 3B). For muscle fiber hypertrophy and atrophy, genes involved with translation like *Eef1a1, Eif4a1, Rpl14, Rpl26*, and *Rpsa* were upregulated in both strains after rotator cuff tear, as were the proteolytic genes *Capn2* and *Nedd8*, while other genes involved with protein degradation and autophagy like *Atg14, Fbxo32, Nedd4, Pik3c3*, and *Ulk1* were downregulated (Figure 3C). Muscle fiber contractility and cytoskeletal organization genes were generally downregulated in B6 and MRL mice after tear, including *Acta1, Actn3, Dag1, Dmd, Dtna, Mypbc2, Myh4, Nf2*, and *Nos1*, while *Actn1, Msn*, and *Myog* were upregulated, and *Myh8* was down in B6 mice and upregulated in MRL mice (Figure 3D). ECM genes were nearly all upregulated in both strains after rotator cuff tear, including *Bgn, Col1a1, Col4a1, Col14a1, Eln, Lox, Loxl1, Mmp2, Mmp13, Mmp14, Thsb4, Timp1, Tnmd*, and *Tnc*, as were the fibroblast markers *S100a4, Scx*, and *Tcf7l2* (Figure 3E). Several growth factors and signaling molecules were also induced in both B6 and MRL mice after rotator cuff tear, such as *Bmp1, Bmp6, Ctgf, Igf1, Pdgfb*, and *Tgfb1*, while *Fgf2, Mstn*, and *Wnt4* were downregulated (Figure 3F). Macrophage cell markers and recruitment genes like *Cd68* and *Cxcl12* were induced (Figure 3F). The muscle innervation marker *Kcnj12* was downregulated, and the denervation marker *Ncam* was upregulated (Figure 3F). Metabolic genes generally showed a downregulation in both strains after rotator cuff tear, including those involved in glycolysis such as *Eno3, Fbp2, Hk2*, as well mitochondrial and oxidative phosphorylation genes *Acads*, *Acss1, Cpt1b, Cpt2, Sdha, Slc25a20*, and *Suclg1* (Figure 3G).

Comparing MRL control to B6 control, 408 transcripts were upregulated by at least 50% and 429 were downregulated by at least 50% (Figure 4A). For tear groups, 497 transcripts were at least 50% upregulated in MRL mice and 402 were downregulated by the same extent (Figure 4D). Many of the genes that were most highly differentially expressed between the strains, either in control or tear groups, are psuedogenes or transcripts that are not yet annotated (Figure 4B-C,E-F). Rotator cuff tear did not affect the expression of many of the most highly differentially expressed genes between strains (Figure 4B-C,E-F). Changes in genes evaluated using RNAseq generally matched observations from qPCR (Supplemental Table S3).

We then sought to explore the effect of rotator cuff tear on changes in the metabolome. PC analysis demonstrated moderate overlap between groups (Figure 5A). For amino acids, arginine, asparagine, aspartate, glutamine, histidine, ornithine, serine, tyrosine, and valine were greater in MRL mice after rotator cuff tear but not in B6 mice, while lysine, phenylalanine, and tryptophan were elevated in both strains (Figure 5B). Glycolytic metabolites were also generally induced, with B6 tear mice uniquely demonstrating elevations in 2-phosphoglycerate/3-phosphoglycerate (2PG/3PG), 6-phosphoglycerate (6PG), erythrose 4-phosphate (E4P), fructose-6-phosphate/glucose-6-phosphate (F6P/G6P), phosphoenolpyruvate (PEP), and ribulose 5-phosphate and xylulose 5-phosphate (R5P/X5P), while acetyl-phosphate (acetyl-P), gluconate, and hexoses were greater in MRL mice (Figure 5C). Creatine, creatinine, and lactate were significantly lower in MRL mice after rotator cuff tear, but not B6 (Figure 5C). Krebs cycle intermediates acetyl Co-A and malate were enriched in B6 mice, while succinate was lower (Figure 5D). Reduced glutathione (GSH), oxidized glutathione (GSSG), and pantothenate were lower in MRL mice after rotator cuff tear (Figure 5D). The nucleotide and nucleoside metabolites AMP, ATP, GDP, GMP, hypoxanthine, and uridine diphosphate-N-acetyl-D-glucosamine (UDP-N-A-D-G) were higher in B6 mice, while GTP, inosine, UDP, and xanthine were uniquely elevated in MRL mice (Figure 5E). CMP, IMP, UMP, and UTP were reduced in MRL mice, and uridine diphosphate-D-glucose (UDP-D-Gluc) was lower in both strains after rotator cuff tear (Figure 5E).

**Figure 5.**
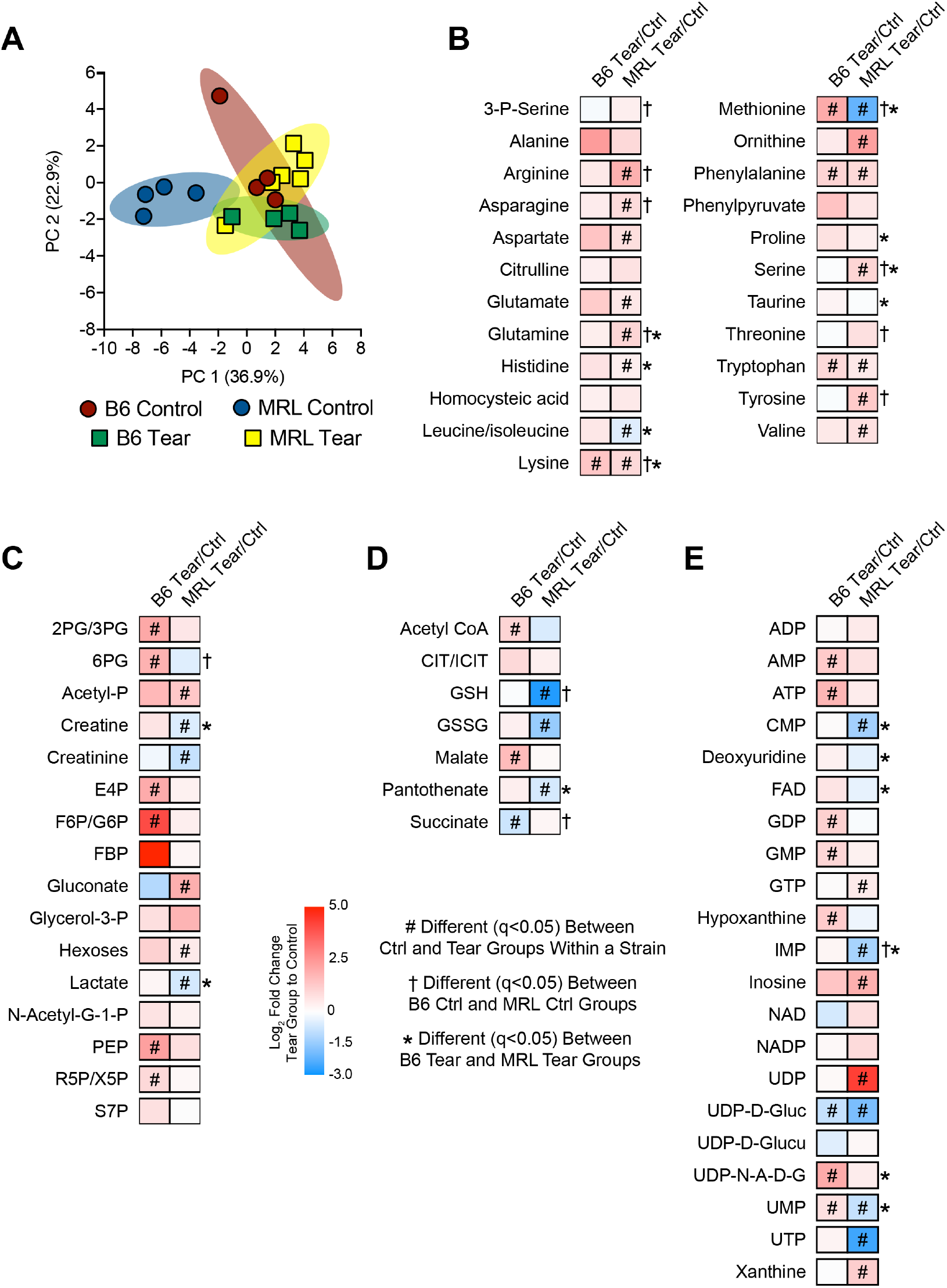
Metabolomics. (A) Principal component (PC) analysis of infraspinatus muscles. Red, B6 control; green, B6 tear; blue, MRL control; yellow, MRL tear. Heatmaps demonstrating the log_2_-fold change of tear groups to controls for both B6 and MRL mice for metabolites related to (B) amino acid, (C) glycolytic, (D) Krebs cycle, and (E) nucleotide and nucleoside metabolism. #, different (q<0.05) between ctrl and tear groups within a strain; †, different (q<0.05) between B6 ctrl and MRL ctrl groups; *, different (q<0.05) between B6 tear and MRL tear groups. N=4 infraspinatus muscles per group except MRL tear which has N=6. Abbreviations: 2PG/3PG, 2-phosphoglycerate and 3-phosphoglycerate; 6PG, 6-phosphoglycerate; CIT/ICIT, citrate and isocitrate; E4P, erythrose 4-phosphate; F6P/G6P, fructose-6-phosphate and glucose-6-phosphate; FBP, fructose bisphosphate; Glycerol-3-P, glycerol-3-phosphate; GSH, reduced glutathione; GSSG, oxidized glutathione; N-Acetyl-G-1-P, N-acetyl-glucosamine-1-phosphate; PEP, phosphoenolpyruvate; R5P/X5P, ribulose 5-phosphate and xylulose 5-phosphate; S7P, sedoheptulose 7-phosphate; UDP-D-Gluc, uridine diphosphate-D-glucose; UDP-D-Glucu, uridine diphosphate-D-glucuronate; and UDP-N-A-D-G, uridine diphosphate-N-acetyl-D-glucosamine.

Finally, we evaluated changes in the lipidome after rotator cuff tear. The PC analysis demonstrated a fair amount of overlap between tear groups, and some divergence in control groups (Figure 6A). The uniquely induced species for B6 mice after rotator cuff tear included cholesterol esters (CE), glucosylceramides (GlcCer), lysophosphatidylethanolamines (LysoPE), and triglycerides (TG), while for MRL mice had elevations in free fatty acids (FFA), phosphatidylcholines (PC), phosphatidylethanolamines (PE), and plasmenyl phosphatidylethanolamines (PlasmenylPE) (Figure 6B). Lysophosphatidylcholines (LysoPC), monoglycerides, and sphingomyelins were induced in both groups (Figure 6B).

**Figure 6.**
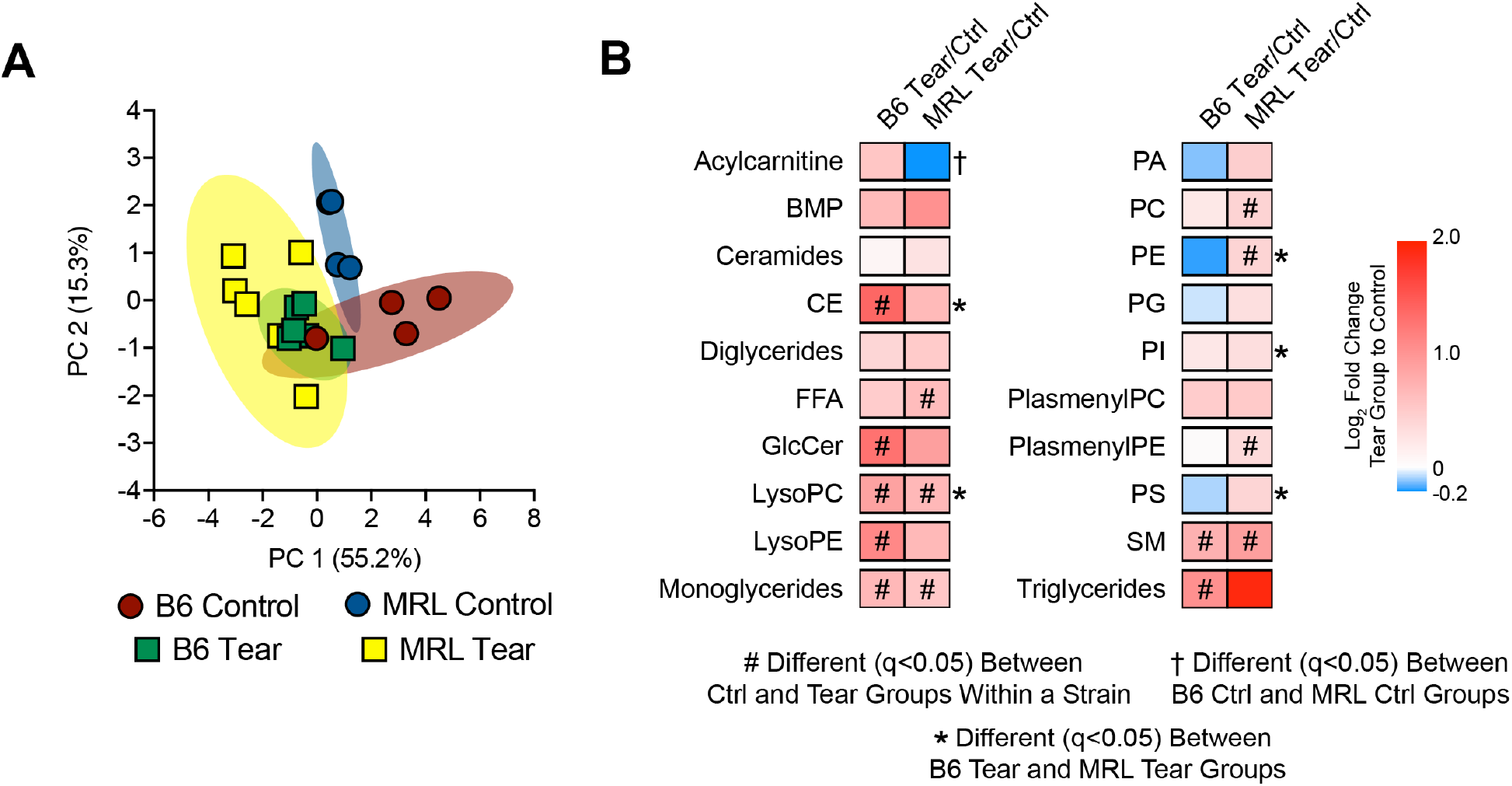
Lipidomics. (A) Principal component (PC) analysis of infraspinatus muscles. Red, B6 control; green, B6 tear; blue, MRL control; yellow, MRL tear. Heatmaps demonstrating the log_2_-fold change of tear groups to controls for both B6 and MRL mice for lipid species. Differences between groups assessed using FDR-corrected t-tests. #, different (q<0.05) between ctrl and tear groups within a strain; †, different (q<0.05) between B6 ctrl and MRL ctrl groups; *, different (q<0.05) between B6 tear and MRL tear groups. N=4 infraspinatus muscles per group for control samples and N=6 infraspinatus muscles per group for tear samples. Abbreviations: BMP, bis(monoacylglycero)phosphates; CE, cholesterol esters; FFA, free fatty acids; GlcCer, glucosylceramides; LysoPC, lysophosphatidylcholines; LysoPE, lysophosphatidylethanolamines; PA, phosphatidic acids; PC, phosphatidylcholines; PE, phosphatidylethanolamines; PG, phosphatidylglycerols; PI, phosphatidylinositols; PlasmenylPC, plasmenyl phosphatidylcholines; PlasmenylPE, plasmenyl phosphatidylethanolamines; PS, phosphatidylserines; and SM, sphingomyelins.

## Discussion

Fatty degeneration, which is commonly observed in patients with rotator cuff tears, limits functional recovery following surgical repair ^3,7^. The MRL mouse strain displays reduced pathological changes and improved recovery in some disease and injury states, ^8-10,12,13^, but not in others ^14,15^. Based on these findings we hypothesized that, compared to B6 mice, MRL mice would have less muscle fiber atrophy and fat accumulation, and be protected against the loss in muscle strength that occurs after rotator cuff tear. While there were widespread differences in gene expression observed between the strains, contrary to our hypothesis we observed no improvement in relative muscle fiber force production or atrophy, and no substantial favorable changes in the metabolome or lipidome between B6 and MRL strains. The general findings from this study indicate that MRL mice are not protected from fatty degeneration after rotator cuff tear.

A pronounced loss of shoulder strength is observed in patients with rotator cuff tears, and often this does not recover even after successful surgical repair and post-operative rehabilitation ^4,7,30,31^. In the current study, control MRL mice had a sF_o_ value that was 52% higher than B6 mice, suggesting that muscle fibers from MRL mice have a greater density of myofibrils per cross-section at baseline. However, while MRL mice had higher sF_o_ values, both strains experienced a 40% reduction in sF_o_ after tear. Using the same surgical model, rats had a 50% reduction in sF_o_ 30 days after rotator cuff tear ^16^. Additionally, patients with a rotator cuff tear have a 29% reduction in sF_o_, and sF_o_ is negatively correlated with patient reported indices of shoulder function ^4^. The findings of reduced force production are consistent with the observed general reduction in contractile and muscle fiber cytoskeletal genes and alterations in muscle regenerative genes in this study, and in previous reports evaluating pathological changes in muscles after rotator cuff tear ^16,21,32-34^. Gene enrichment analysis predicted an increase in IGF1 signaling, which is consistent with observed at the transcript and protein level in a rat model of rotator cuff tears ^16^. In resistance exercise training programs, IGF1 is known to induce skeletal muscle hypertrophy and increased force production ^35,36^. However, in the context of rotator cuff tears, even though IGF1 signaling is active ^16^ this does not appear sufficient to protect against a loss in force production after injury.

Fibrosis is a common finding in injured rotator cuff muscles ^21,33,34^. In the current study, there was a general upregulation in the fibrillar collagen type I (*Col1a1* and *Col1a2*), the basement membrane collagen type IV (*Col4a1*), elastin (*Eln*), as well as the proteoglycans biglycan (*Bgn*), thrombospondin 4 (*Thsb4*), tenomodulin (*Tnmd*), and tenascin C (*Tnc*) after injury. Fibroblasts synthesize and remodel the muscle ECM ^37^, and markers of fibroblasts such as *S100a4, Scx*, and *Tcf7l2* were also upregulated and not different between B6 and MRL mice after rotator cuff tear. Based on findings that γ-sarcoglycan/MRL mice had lower hydroxyproline levels and improvements in fibrosis compared to γ-sarcoglycan mice ^11^ we anticipated that MRL mice would have less collagen and fibroblast marker gene expression after injury. However, our results indicate no difference in markers of fibrosis between B6 and MRL mice after rotator cuff tear.

Changes in the muscle metabolome and lipidome also have important functional consequences in the context of rotator cuff tears. Using a rat model of rotator cuff tears, we demonstrated that mitochondrial dysfunction leads to the progressive increase in lipid accumulation that occurs after rotator cuff tear ^16^. We did not directly measure mitochondrial function in the current study, but gene enrichment analysis predicted mitochondrial dysfunction was present in both B6 and MRL groups after rotator cuff tear, and there was a downregulation in key genes involved in β-oxidation of lipids in mitochondria and aerobic metabolism. These genes include as acyl-CoA synthetases (*Acads*) and dehydrogenase (*Acss1*), carnitine palmitoyl transferases (*Cpt1b* and *Cpt2*), succinate dehydogenase (*Sdha*), carnitine acylcarnitine translocase (*Slc25a20*), and succinyl CoA synthetase (*Suclg1*). These findings are in agreement with observation in rats ^16^, but there was some divergence in glycolytic metabolism differences between rats, and B6 and MRL mouse muscles after rotator cuff tear. There was a marked induction in glycolytic metabolites in rats, and B6 muscles showed a similar increase in several of the same metabolites. However, MRL mice only had slight changes in glycolytic metabolites but substantial increases in several free amino acids. These findings may indicate strain-specific differences in substrate metabolism, with B6 mice possibly relying on glycolytic metabolism, while MRL mice might shift to greater amino acid and ketone metabolism after rotator cuff tear. An increased reliance on amino acid metabolism in the MRL genetic background is supported by an observed elevation in histidine ammonia-lyase (*Hal*), which is a critical enzyme involved in histidine and glutamate metabolism ^38^ and is one of the most highly upregulated genes in MRL mice both before and after rotator cuff tear.

There were also differences in lipid species between B6 and MRL mice. At 30 days after a rotator cuff tear, rats demonstrated an increase in triglycerides, diglycerides, FFA, CE, LysoPC, PC, PE, PG, PI, PS, and SM, with no change in monoglycerides, LysoPE, or PlasmenylPE ^16^. In agreement with these findings, B6 mice had an elevation in triglycerides and CE, while MRL mice had increased FFA, PC, and PE, and both strains had higher levels of LysoPC and SM. Different from rats with rotator cuff tears, B6 mice demonstrated increased LysoPE, and MRL mice had higher PlasmenylPE, with both having elevated monoglycerides. Taken together, these results indicate that there are strain- and species-specific changes that occur in the metabolome and lipidome after rotator cuff tear. The differences in response between B6 and MRL mice, however, do not impact functional changes in the protection against weakness after rotator cuff tear. There may also be intrinsic differences in lipid metabolism between strains that explain the divergent response to a rotator cuff tear. For example, acetyl-CoA acyltransferase (*Acaa1b*), which is a critical enzyme involved in non-mitochondrial peroxisomal oxidation of lipids ^39^, was one of the most highly upregulated genes in MRL mice prior to injury.

There are several limitations to this study. We only included male mice, and it is possible that the MRL background would confer greater protection against fatty degeneration in female mice. Due to their quadripedal gait, the mechanical loading demands of the rotator cuff of mice is different from humans. We did not repair torn rotator cuffs in this study, and it is possible that there are strain-specific effects on rotator cuff healing that occur independent of the development of fatty degeneration. A single time point was evaluated in this study, and there could be differences between B6 and MRL mice at later time points. We only evaluated 3 month old mice, and there is known to be an aging-associated exacerbation in pathological changes in rotator cuff tears ^40^. Despite these limitations, we think this report provides important insight into the biological changes in muscle function after rotator cuff tear.

Rotator cuff tears can result in profound pain and disability that persist despite a technically successful surgical repair ^1,7,41^. Although surgical techniques and rehabilitation programs have evolved to improve rotator cuff healing, our ability to treat the extensive muscle atrophy and fatty degeneration that occur as a result of the tear is limited ^3^. While there were numerous differences in the transcriptome, metabolome, and lipidome between B6 and MRL mice, this did not lead to any functional differences in relative muscle fiber force production between strains. The results provide further support to the notion that the fat which accumulates after rotator cuff likely occurs due to deficits in lipid oxidation, leading to a pro-inflammatory and lipotoxic state that impairs regeneration. Although the MRL background did not convey functional protection against pathological changes in muscles after rotator cuff tear, several studies have shown that MRL mice have superior tendon healing properties compared to B6 mice ^12,13,42^. If these findings of improved intratendon healing translate to enhanced bone-tendon insertion site healing, gaining a greater understanding of the biological pathways that convey improved tendon healing in MRL mice may lead to new therapies for rotator cuff tears, although it is not clear that these would directly impact muscle regeneration.

## Acknowledgements

This work was funded by NIH grant R01-AR063649. The University of Michigan Metabolomics Core is supported by NIH grant U24-DK097153. We would like to acknowledge technical assistance from Dr. David Oliver and Mr. Damonie Salmon at the Hospital for Special Surgery.

## Author Contribution Statement

JRT, SW, JSD, SAR, and CLM designed research; JRT, ANP, SW, ACK, JBS, and SN acquired data; JRT, APN, and CLM analyzed and interpreted data; JRT and CLM drafted the paper or revised the paper critically; all authors approved of the submitted and final version.

