## Supplemental Table S1 for "The MRL/MpJ Mouse Strain Is Not Protected From Muscle Atrophy And Weakness After Rotator Cuff Tear"

***Supplemental Table S1. Metabolomics data.*** Data presented as mean relative peak intensity (RPI) for each group. N=4 muscles per group except MRL tear which is N=6.

| **Analyte** | **B6 No Tear (Mean RPI)** | **B6 Tear (Mean RPI)** | **MRL No Tear (Mean RPI)** | **MRL Tear (Mean RPI)** |
|---|---|---|---|---|
| 3-P-Serine | 6099817.528 | 5639293.179 | 2556748.822 | 3318021.1 |
| Alanine | 54064192.83 | 268825273.7 | 69868303.12 | 129502274.7 |
| Arginine | 137877470.1 | 196248439.9 | 54463536.83 | 198760711.7 |
| Asparagine | 91328174.54 | 124093166 | 42365209.73 | 75828550.64 |
| Aspartate | 137927722.7 | 360725736.2 | 140279140.6 | 236017797.2 |
| Citrulline | 4605107.706 | 5895217.341 | 2467358.357 | 3938143.851 |
| Glutamate | 371518993.9 | 859983149.6 | 457190345.5 | 667488778.5 |
| Glutamine | 1062818524 | 1393484154 | 443938880.6 | 873749719.3 |
| Histidine | 856427601.1 | 1346868867 | 579046138.7 | 751798403.1 |
| Homocysteic acid | 711486896.8 | 906637853.8 | 382384374.1 | 539878976 |
| Leucine/isoleucine | 9449464927 | 13680072249 | 11362828026 | 8182521399 |
| Lysine | 293767193.6 | 748650222.1 | 185642261.3 | 342931753.2 |
| Methionine | 79575638.54 | 316462768.1 | 305796783.7 | 69850968.43 |
| Ornithine | 16905954.13 | 23284281.93 | 6109257.641 | 28472805.9 |
| Phenylalanine | 830342974.8 | 1688157620 | 703718572.3 | 1320137892 |
| Phenylpyruvate | 965017.5116 | 2564196.803 | 972546.8347 | 1423484.879 |
| Proline | 295758500.3 | 477599585.5 | 260308935.6 | 339338648.8 |
| Serine | 154608055.6 | 149408104.2 | 46035519.81 | 91323009.2 |
| Taurine | 43052099656 | 50892914521 | 40214789223 | 38664810058 |
| Threonine | 115946295 | 111868922.5 | 39429667.14 | 64959229.25 |
| Tryptophan | 160311565.8 | 297082082.6 | 149170362 | 214236623.6 |
| Tyrosine | 169831045.5 | 162419327.1 | 59937711.27 | 140806129 |
| Valine | 748179751.3 | 1065896829 | 602139149.1 | 994571407.8 |
| 2PG/3PG | 232.2756662 | 954.744949 | 952.7318603 | 1403.997565 |
| 6PG | 3142707.101 | 11064276.73 | 13330682.96 | 9631408.694 |
| Acetylphosphate | 37110511.23 | 117312444 | 44972509.9 | 111374403.9 |
| Creatine | 8712432382 | 13399134240 | 11516542790 | 8112161513 |
| Creatinine | 159565161 | 139759374.9 | 185973220.9 | 104382398.1 |
| E4P | 573.7524431 | 2251.125225 | 1439.832079 | 1789.623381 |
| F6P/G6P | 157.7648985 | 2568.478078 | 1525.254888 | 1984.945356 |
| FBP | 33.11013733 | 1056.154032 | 852.8904116 | 975.8354888 |
| Gluconate | 130853392.7 | 61129470.94 | 41452977.22 | 153431116.4 |
| Glycerol-3-P | 5.858095039 | 9.738278895 | 6.700296076 | 22.91053592 |
| Hexoses | 1861641605 | 3906992136 | 2339253484 | 3297550314 |
| LAC | 11178301409 | 12726378812 | 13932497055 | 8992176764 |
| N-Acetyl-G-1-P | 17931194.9 | 27850112.65 | 18456085.46 | 22879422.32 |
| PEP | 5.802680436 | 27.15128849 | 19.62761652 | 32.65058916 |
| R5P/X5P | 500.9604315 | 919.6123099 | 609.1529602 | 677.6625669 |
| s7P | 19.94093628 | 32.86315619 | 20.47770677 | 20.68095772 |
| Acetyl CoA | 200741.9661 | 408011.4907 | 620126.7101 | 421402.1197 |
| CIT/ICIT | 54.6016993 | 100.6909501 | 110.8860386 | 141.3571503 |
| GSH | 189409405 | 182663266.1 | 878319111.6 | 125302546.8 |
| GSSG | 168667213.7 | 212923815.9 | 486959927.9 | 174277450.5 |
| MAL | 266.4764932 | 806.8445941 | 491.6064978 | 532.466974 |
| Pantothenate | 973368999.3 | 1257030765 | 1252542342 | 812125264.6 |
| SUC | 66088946.3 | 39344757.27 | 29246680.33 | 33369250.34 |
| ADP | 163.4808247 | 174.4299852 | 181.9475971 | 253.6606607 |
| AMP | 25.50539729 | 60.07541176 | 47.0904081 | 75.03848245 |
| ATP | 30.21469082 | 100.852698 | 80.43177048 | 107.6313759 |
| CMP | 68364984.31 | 72168982.22 | 115876549.4 | 45900938.38 |
| Deoxyuridine | 115054107.4 | 144880893.4 | 113908455.8 | 87119220.64 |

| | | | | |
|---|---|---|---|---|
| FAD | 15707034.9 | 24074539.98 | 18201371.37 | 14345777.76 |
| GDP | 3115610.844 | 6638319.596 | 5306116.657 | 4976166.784 |
| GMP | 59.06966066 | 114.4398934 | 88.16510691 | 109.435388 |
| GTP | 11.94530069 | 12.56487382 | 8.351824521 | 11.59204214 |
| Hypoxanthine | 66445741.89 | 152885019.3 | 115040889.2 | 97302296.74 |
| IMP | 9647454334 | 11119726048 | 18089685355 | 7484436524 |
| Inosine | 646168404.3 | 1665709696 | 441321243.8 | 1612260086 |
| NAD | 7.497179328 | 4.842779829 | 3.2137154 | 5.44127189 |
| NADP | 8.645017213 | 9.456127568 | 9.126281867 | 16.29804061 |
| UDP | 12481269.98 | 13524361.38 | 954569.8304 | 18566721.38 |
| UDP-D-glucose | 34903380.13 | 19191967.95 | 47543991.86 | 13078576.09 |
| UDP-D-glucuronate | 43419384.62 | 31470362.22 | 24967650.9 | 27605275.43 |
| UDP-N-A-D-G | 54578069.39 | 214698481.2 | 72838323.86 | 97977976.41 |
| UMP | 432672163.3 | 704076639.8 | 686662837.9 | 382183642.2 |
| UTP | 802695.7553 | 950099.7329 | 5095082.865 | 872294.5527 |
| Xanthine | 1988715164 | 1942809172 | 1157809249 | 2582672194 |
