## Supplemental Table S2 for "The MRL/MpJ Mouse Strain Is Not Protected From Muscle Atrophy And Weakness After Rotator Cuff Tear"

***Supplemental Table S2. Lipidomics data.*** Data presented as mean relative peak intensity (RPI) for each group. N=4 muscles per group for no tear and N=6 muscles per group for tear.

| **Analyte** | **B6 No Tear (Mean RPI)** | **B6 Tear (Mean RPI)** | **MRL No Tear (Mean RPI)** | **MRL Tear (Mean RPI)** |
|---|---|---|---|---|
| 16:0 Cholesteryl ester | 904.14 | 1743.84 | 749.14 | 948.98 |
| 16:1 Cholesteryl ester | 508.73 | 1143.47 | 994.67 | 1843.80 |
| 18:1 Cholesteryl ester | 1405.32 | 1847.99 | 3495.33 | 4543.38 |
| 18:2 Cholesteryl ester | 9111.10 | 31288.47 | 27699.94 | 75017.29 |
| 18:3 Cholesteryl ester | 3580.27 | 1919.75 | 438.39 | 5074.87 |
| 20:3 Cholesteryl ester | 1158.97 | 6011.22 | 4606.25 | 29367.49 |
| 20:4 Cholesteryl ester | 3902.92 | 3529.03 | 20553.60 | 12288.43 |
| 20:5 Cholesteryl ester | 1943.12 | 1568.79 | 2266.23 | 3375.51 |
| 22:4 Cholesteryl ester | 1249.34 | 2312.78 | 468.40 | 3927.40 |
| 22:6 Cholesteryl ester | 2854.54 | 3612.22 | 13665.83 | 5690.91 |
| Acylcarnitine 14:0 | 41103.40 | 36095.89 | 66302.12 | 59913.67 |
| Acylcarnitine 16:0 | 321445.38 | 411749.35 | 1084433.25 | 602228.83 |
| Acylcarnitine 16:1 | 35658.97 | 117786.30 | 91936.57 | 358656.28 |
| Acylcarnitine 16:2 | 10929.23 | 10971.36 | 18153.85 | 14613.70 |
| Acylcarnitine 17:0 | 18602.33 | 19226.24 | 24347.20 | 39975.33 |
| Acylcarnitine 18:0 | 111664.02 | 102353.24 | 248638.26 | 171083.42 |
| Acylcarnitine 18:1 | 188360.09 | 221367.37 | 532938.53 | 401475.23 |
| Acylcarnitine 18:2 | 68548.95 | 243542.43 | 203799.84 | 484058.00 |
| Acylcarnitine 20:1 | 9913.85 | 11313.62 | 43056.24 | 28352.22 |
| Acylcarnitine 24:0 | 2049.27 | 3001.59 | 3451.18 | 4382.02 |
| BMP 36:4 | 6615.31 | 18894.76 | 9789.41 | 18390.27 |
| BMP 38:6 | 7094.41 | 10829.06 | 5553.04 | 14373.30 |
| BMP 38:7 | 7642.70 | 6779.35 | 5706.06 | 9963.35 |
| BMP 40:7 | 8603.43 | 12152.91 | 7018.98 | 17589.70 |
| BMP 40:8 | 44923.26 | 72198.60 | 28007.92 | 91133.18 |
| BMP 44:12 | 9309.76 | 10881.00 | 9503.99 | 14689.35 |
| Cer[AS] 34:1 | 6757.39 | 8732.78 | 5076.71 | 10477.02 |
| Cer[AS] 40:1 | 6188.31 | 61573.98 | 14457.49 | 49901.58 |
| Cer[AS] 42:1 | 15313.31 | 131011.44 | 17470.73 | 82306.44 |
| Cer[EODS] 51:0 | 14049.08 | 9433.71 | 12679.73 | 14643.44 |
| Cer[EODS] 53:0 | 9360.33 | 6057.90 | 7827.83 | 7240.22 |
| Cer[NDS] 36:0 | 513901.43 | 373085.39 | 464099.17 | 505316.29 |
| Cer[NDS] 40:0 | 4303.33 | 29904.72 | 5475.18 | 35898.82 |
| Cer[NDS] 41:0 | 9058.97 | 8914.33 | 7499.49 | 12632.51 |
| Cer[NDS] 42:0 | 10941.74 | 21756.72 | 11168.30 | 16725.35 |
| Cer[NP] 35:1 | 40057.41 | 33458.48 | 36033.59 | 56324.68 |
| Cer[NP] 36:0 | 53072.83 | 42780.35 | 53981.88 | 56952.65 |
| Cer[NP] 36:1 | 63376.91 | 30375.92 | 55728.64 | 53859.83 |
| Cer[NP] 40:0 | 27812.23 | 78764.04 | 32992.92 | 67726.47 |
| Cer[NP] 41:0 | 42182.52 | 41440.34 | 32811.67 | 47365.00 |
| Cer[NP] 42:0 | 51077.05 | 121666.06 | 43106.64 | 93413.98 |
| Cer[NS] 32:1 | 5671.08 | 6286.41 | 4180.01 | 9151.43 |
| Cer[NS] 33:1 | 31477.83 | 32995.47 | 23240.64 | 46145.54 |
| Cer[NS] 34:1 | 169083.25 | 315480.69 | 148994.28 | 404191.76 |
| Cer[NS] 34:2 | 7688.75 | 25439.97 | 9315.67 | 25111.70 |
| Cer[NS] 36:1 | 6094270.73 | 4976370.16 | 5542584.61 | 5073651.89 |
| Cer[NS] 36:2 | 208747.76 | 226923.76 | 205316.85 | 244723.51 |
| Cer[NS] 36:3 | 6960.26 | 7344.50 | 10235.42 | 10123.06 |
| Cer[NS] 37:1 | 608784.36 | 381506.97 | 523595.37 | 347099.24 |
| Cer[NS] 37:2 | 32666.54 | 28074.40 | 31857.67 | 26014.25 |
| Cer[NS] 38:1 | 246774.74 | 218258.03 | 206992.09 | 288008.74 |

| | | | | |
|---|---|---|---|---|
| Cer[NS] 38:2 | 33359.49 | 50603.31 | 38569.56 | 67788.32 |
| Cer[NS] 39:1 | 12927.18 | 14589.86 | 5568.57 | 19246.28 |
| Cer[NS] 39:2 | 6368.66 | 15966.74 | 6894.50 | 18002.32 |
| Cer[NS] 40:1 | 979293.49 | 1129071.21 | 730621.24 | 1457784.20 |
| Cer[NS] 40:2 | 88404.73 | 190070.37 | 83681.06 | 252716.34 |
| Cer[NS] 41:1 | 813387.00 | 974234.78 | 624660.07 | 928050.43 |
| Cer[NS] 41:2 | 12010.85 | 12627.25 | 8293.55 | 15922.96 |
| Cer[NS] 42:1 | 898263.56 | 1805994.25 | 897079.90 | 1654940.30 |
| Cer[NS] 42:2 | 112710.99 | 179036.41 | 84344.46 | 172590.68 |
| Cer[NS] 42:3 | 269683.47 | 551423.24 | 176418.60 | 553525.95 |
| Cer[NS] 42:4 | 46504.62 | 42250.98 | 50176.46 | 70924.05 |
| Cer[NS] 43:1 | 56316.19 | 81539.21 | 60693.95 | 80599.43 |
| Cer[NS] 43:2 | 2990.70 | 5806.68 | 4340.72 | 7227.46 |
| Cer[NS] 44:2 | 3404.59 | 5221.93 | 2550.50 | 5516.48 |
| DG 30:0 | 46726.50 | 52874.00 | 63479.73 | 88297.77 |
| DG 30:1 | 7052.90 | 11001.66 | 10477.66 | 21924.02 |
| DG 31:0 | 8539.55 | 11562.43 | 9326.87 | 11661.42 |
| DG 32:0 | 223826.19 | 274412.72 | 347745.16 | 333298.55 |
| DG 32:1 | 184410.50 | 327216.81 | 280387.96 | 486062.33 |
| DG 32:2 | 82296.81 | 125694.38 | 81718.36 | 195637.38 |
| DG 32:3 | 15327.45 | 24317.90 | 9091.06 | 45001.62 |
| DG 33:0 | 9798.59 | 11226.10 | 10757.81 | 12034.10 |
| DG 33:1 | 22410.19 | 19262.55 | 22406.14 | 23849.45 |
| DG 33:2 | 19131.63 | 25251.51 | 18071.52 | 32390.23 |
| DG 34:0 | 104334.69 | 100951.40 | 102432.87 | 132252.63 |
| DG 34:1 | 952797.28 | 762131.08 | 847172.75 | 1150590.52 |
| DG 34:2 | 949033.59 | 1610515.70 | 1082999.84 | 2047382.82 |
| DG 34:3 | 205878.76 | 229908.80 | 150886.56 | 341906.33 |
| DG 34:4 | 9776.83 | 43731.48 | 11365.43 | 72107.28 |
| DG 35:0 | 6201.95 | 8775.76 | 6897.80 | 7939.28 |
| DG 35:1 | 16033.48 | 21129.70 | 19471.80 | 24559.08 |
| DG 35:2 | 13032.45 | 62653.67 | 13821.24 | 72993.14 |
| DG 35:3 | 9804.50 | 50590.30 | 11986.99 | 71427.84 |
| DG 36:0 | 72252.53 | 66163.47 | 81414.33 | 83265.32 |
| DG 36:1 | 99490.97 | 141550.39 | 136384.19 | 195843.66 |
| DG 36:2 | 394620.65 | 691721.44 | 404597.68 | 847391.11 |
| DG 36:3 | 736191.52 | 986516.81 | 658705.96 | 1215006.40 |
| DG 36:4 | 557074.58 | 645773.15 | 453819.71 | 769056.99 |
| DG 36:5 | 62361.03 | 89234.46 | 44460.17 | 174289.58 |
| DG 36:6 | 12070.36 | 15029.06 | 11400.65 | 23974.96 |
| DG 37:1 | 11145.67 | 17540.96 | 8072.63 | 22392.21 |
| DG 37:2 | 20145.25 | 36925.30 | 16518.59 | 37734.04 |
| DG 37:3 | 8369.97 | 33560.44 | 6861.01 | 27095.24 |
| DG 38:0 | 1110520.70 | 960628.89 | 1078761.43 | 1434747.56 |
| DG 38:1 | 18181.23 | 20225.99 | 21790.11 | 27406.73 |
| DG 38:2 | 53605.96 | 57843.73 | 61238.90 | 72021.51 |
| DG 38:3 | 105295.27 | 153653.11 | 115827.19 | 166615.17 |
| DG 38:4 | 124821.77 | 744722.83 | 144776.38 | 892232.49 |
| DG 38:5 | 158051.57 | 143435.73 | 211992.91 | 210628.98 |
| DG 38:6 | 307049.93 | 236023.65 | 365451.21 | 312726.70 |
| DG 38:7 | 37946.34 | 30486.21 | 22342.30 | 61769.61 |
| DG 40:0 | 2863.29 | 3506.65 | 2963.92 | 4306.30 |
| DG 40:3 | 7851.12 | 8860.79 | 9719.89 | 9672.03 |
| DG 40:4 | 11816.24 | 13283.42 | 14143.61 | 20172.49 |
| DG 40:5 | 44951.86 | 50742.91 | 60596.36 | 65428.63 |
| DG 40:6 | 99873.49 | 97163.80 | 109588.86 | 130201.25 |

| | | | | |
|---|---|---|---|---|
| DG 40:7 | 150330.60 | 132476.40 | 178834.38 | 136552.05 |
| DG 40:8 | 52091.09 | 168120.05 | 51779.53 | 188748.40 |
| DG 41:6 | 4297.19 | 4739.24 | 4415.31 | 4932.71 |
| DG 42:6 | 10496.11 | 9090.71 | 17457.13 | 11951.07 |
| DG 42:7 | 13761.41 | 12906.97 | 23781.60 | 14698.21 |
| DG 42:8 | 10729.75 | 9456.32 | 15159.57 | 11012.21 |
| DG 42:9 | 8650.17 | 7000.12 | 14514.56 | 10468.11 |
| FFA(16:0) | 45731054.63 | 59093202.30 | 48923397.73 | 77842359.18 |
| FFA(18:0) | 56638999.68 | 65642884.96 | 59267670.45 | 87894524.55 |
| FFA(18:1) | 12009591.96 | 23034746.44 | 13970763.35 | 28951530.10 |
| FFA(18:2) | 11545506.41 | 27134112.57 | 13917286.51 | 28138735.06 |
| FFA(20:0) | 2454267.99 | 1234380.63 | 1636506.97 | 2035818.24 |
| FFA(20:1) | 846902.92 | 1793172.49 | 1269358.41 | 1927960.33 |
| FFA(20:2) | 315157.13 | 761160.32 | 443119.27 | 817536.22 |
| FFA(20:4) Arachidonic acic | 1821152.94 | 4773404.60 | 2705884.87 | 7537516.30 |
| FFA(22:0) | 692879.63 | 727805.59 | 672608.72 | 980799.56 |
| FFA(22:1) | 305919.55 | 446480.08 | 353429.30 | 553169.84 |
| FFA(22:2) | 46447.95 | 62165.08 | 47815.96 | 86128.34 |
| FFA(22:3) | 38843.51 | 67148.23 | 62632.43 | 108326.89 |
| FFA(24:0) | 187812.06 | 185547.92 | 165819.92 | 260446.25 |
| FFA(24:1) | 115373.78 | 220942.68 | 84911.88 | 185258.22 |
| FFA(24:2) | 20945.85 | 50817.81 | 17674.69 | 64692.69 |
| FFA(24:3) | 3163.28 | 11868.06 | 2778.13 | 14992.30 |
| GlcCer[NS] 34:1 | 11047.59 | 33884.70 | 3867.31 | 25277.76 |
| GlcCer[NS] 36:1 | 8001.48 | 9415.74 | 7056.73 | 14093.75 |
| GlcCer[NS] 38:2 | 9662.83 | 21199.93 | 10345.41 | 25719.92 |
| GlcCer[NS] 40:1 | 51659.65 | 155489.62 | 75648.49 | 172745.07 |
| GlcCer[NS] 40:2 | 25654.98 | 137860.54 | 30394.12 | 197199.01 |
| GlcCer[NS] 41:1 | 58787.51 | 146937.21 | 61317.08 | 145225.63 |
| GlcCer[NS] 41:2 | 13101.98 | 27394.85 | 15382.47 | 27962.65 |
| GlcCer[NS] 42:1 | 327230.77 | 738243.69 | 453448.20 | 782356.52 |
| GlcCer[NS] 42:2 | 58165.67 | 109605.53 | 72517.23 | 147450.66 |
| GlcCer[NS] 43:1 | 4527.57 | 10044.76 | 2650.41 | 10712.78 |
| LysoPC 14:0 | 7745.98 | 10738.42 | 5136.51 | 8957.81 |
| LysoPC 15:0 | 5302.13 | 9173.00 | 7290.88 | 12594.08 |
| LysoPC 16:0 | 649991.07 | 1169186.73 | 1111510.73 | 1836740.94 |
| LysoPC 16:1 | 21260.19 | 37879.11 | 33316.65 | 52558.07 |
| LysoPC 17:1 | 3375.67 | 6602.40 | 5391.71 | 8192.74 |
| LysoPC 18:0 | 549930.55 | 1053943.39 | 885324.96 | 1847423.59 |
| LysoPC 18:1 | 90403.84 | 235266.86 | 185741.08 | 394325.84 |
| LysoPC 18:2 | 152635.48 | 194241.14 | 182091.72 | 306983.03 |
| LysoPC 18:3 | 13002.55 | 60680.70 | 17111.65 | 72159.93 |
| LysoPC 19:0 | 31081.34 | 44661.23 | 42230.77 | 72712.88 |
| LysoPC 20:0 | 32816.18 | 25341.58 | 29429.18 | 48969.28 |
| LysoPC 20:1 | 6531.48 | 15227.70 | 12550.74 | 30101.19 |
| LysoPC 20:2 | 7916.30 | 9659.90 | 12581.25 | 13906.35 |
| LysoPC 20:3 | 16666.90 | 50346.30 | 22844.88 | 89516.37 |
| LysoPC 20:4 | 61193.79 | 80655.10 | 80732.96 | 159535.33 |
| LysoPC 20:5 | 23562.67 | 59890.01 | 35236.50 | 75220.34 |
| LysoPC 22:0 | 3816.12 | 34306.34 | 6103.86 | 41661.95 |
| LysoPC 22:4 | 1668.12 | 17858.33 | 3506.58 | 26092.64 |
| LysoPC 22:5 | 39874.05 | 78274.86 | 76324.69 | 108963.82 |
| LysoPC 22:6 | 300129.66 | 511277.16 | 573097.65 | 708115.38 |
| LysoPC 24:0 | 9627.19 | 22814.87 | 12034.65 | 22858.95 |
| LysoPC 24:1 | 1265.79 | 3709.01 | 2167.19 | 5315.91 |
| LysoPE 16:0 | 42137.05 | 36845.43 | 35281.34 | 62036.36 |

| | | | | |
|---|---|---|---|---|
| LysoPE 17:0 | 14375.51 | 23863.92 | 15262.02 | 29284.82 |
| LysoPE 18:0 | 106584.50 | 165194.62 | 150710.82 | 265789.50 |
| LysoPE 18:1 | 23488.07 | 55872.80 | 39349.19 | 96349.50 |
| LysoPE 18:2 | 22906.36 | 51901.45 | 25036.40 | 82383.38 |
| LysoPE 20:3 | 4007.57 | 12863.92 | 5304.32 | 13004.71 |
| LysoPE 20:4 | 27437.92 | 89313.60 | 36699.10 | 122279.69 |
| LysoPE 22:4 | 5935.59 | 31770.36 | 14268.52 | 47023.12 |
| LysoPE 22:5 | 12605.82 | 93254.44 | 16574.45 | 102668.85 |
| LysoPE 22:6 | 157912.85 | 333389.51 | 284976.53 | 324017.40 |
| LysoPG 17:0 | 3333.05 | 6224.26 | 2066.17 | 3277.29 |
| MGDG 32:0 | 22090.94 | 31994.31 | 25407.01 | 39464.25 |
| MGDG 34:1 | 5666.70 | 12067.36 | 10143.96 | 16116.30 |
| PA 34:1 | 21296.08 | 16258.96 | 14033.12 | 24020.75 |
| PA 34:2 | 8276.94 | 7714.74 | 5749.34 | 14405.69 |
| PA 36:4 | 5927.05 | 8939.41 | 6568.15 | 10014.44 |
| PA 38:4 | 18025.47 | 13305.40 | 15240.47 | 19276.84 |
| PA 38:6 | 18460.37 | 18047.44 | 14608.81 | 17092.36 |
| PA 40:7 | 26635.26 | 26650.15 | 23349.34 | 29387.06 |
| PA 40:8 | 14534.51 | 9868.12 | 14112.37 | 9647.90 |
| PC 26:1 | 12387.41 | 3304.23 | 1879.44 | 8249.60 |
| PC 28:0 | 106210.78 | 241495.67 | 10514.78 | 34900.72 |
| PC 29:0 | 65578.40 | 69378.52 | 13629.28 | 22020.77 |
| PC 30:0 | 1440603.58 | 1537929.79 | 444813.93 | 693987.90 |
| PC 30:1 | 96498.26 | 161908.68 | 35788.76 | 58323.13 |
| PC 30:2 | 29394.67 | 56090.34 | 7919.08 | 13537.26 |
| PC 31:0 | 148387.51 | 87296.59 | 91570.85 | 100982.89 |
| PC 31:1 | 162345.24 | 133605.47 | 81692.38 | 89901.00 |
| PC 31:2 | 25393.96 | 132957.47 | 10706.30 | 70359.86 |
| PC 32:0 | 6365245.42 | 10668836.24 | 9262173.42 | 11594097.29 |
| PC 32:1 | 1820247.85 | 1606486.41 | 1316569.23 | 2051499.04 |
| PC 32:2 | 2096250.77 | 2608098.86 | 984108.66 | 1264082.66 |
| PC 32:3 | 220046.56 | 304432.93 | 86070.43 | 103150.45 |
| PC 32:4 | 23639.94 | 46222.25 | 16682.76 | 22590.65 |
| PC 33:0 | 140565.93 | 186832.64 | 162786.69 | 224621.54 |
| PC 33:1 | 140528.63 | 174134.70 | 128359.75 | 119159.13 |
| PC 33:2 | 247834.48 | 928018.46 | 174185.60 | 654345.69 |
| PC 33:3 | 111623.81 | 105508.21 | 84812.08 | 83975.18 |
| PC 34:1 | 12567198.07 | 13931639.24 | 12808452.69 | 18640372.72 |
| PC 34:2 | 18821603.37 | 21796628.04 | 16473951.29 | 22067889.49 |
| PC 34:3 | 589010.99 | 806029.71 | 660952.04 | 659054.39 |
| PC 34:4 | 845857.90 | 1009928.58 | 574251.52 | 597389.16 |
| PC 34:5 | 34017.59 | 44984.73 | 30288.56 | 34933.17 |
| PC 34:6 | 21150.91 | 26140.13 | 13177.65 | 25722.24 |
| PC 35:0 | 53881.67 | 217240.98 | 48477.78 | 261930.16 |
| PC 35:1 | 318047.84 | 449959.90 | 342597.04 | 459682.58 |
| PC 35:2 | 78612.16 | 139321.10 | 82157.91 | 110107.48 |
| PC 35:3 | 44858.69 | 66990.65 | 34455.38 | 47057.19 |
| PC 35:4 | 57557.52 | 66404.07 | 57892.84 | 72983.70 |
| PC 35:5 | 87443.40 | 87391.39 | 78234.33 | 97804.53 |
| PC 36:0 | 68812.57 | 82424.28 | 74009.57 | 110700.28 |
| PC 36:1 | 3129542.08 | 7420917.77 | 4206939.97 | 6436181.66 |
| PC 36:2 | 7234349.29 | 9952180.87 | 5618187.48 | 12222193.64 |
| PC 36:3 | 1457887.78 | 1829042.46 | 1394663.94 | 2090282.67 |
| PC 36:4 | 2790004.36 | 2670411.96 | 3007342.21 | 3959911.44 |
| PC 36:5 | 3168658.22 | 3386031.44 | 5414044.98 | 5660242.08 |
| PC 36:6 | 2353047.29 | 2663267.34 | 804634.97 | 926113.06 |

| | | | | |
|---|---|---|---|---|
| PC 36:7 | 30285.96 | 87938.91 | 63903.48 | 79284.98 |
| PC 36:8 | 23719.87 | 56705.48 | 14270.76 | 32336.35 |
| PC 37:1 | 74149.69 | 92814.77 | 73657.38 | 125976.36 |
| PC 37:2 | 160139.91 | 990221.36 | 138712.77 | 916795.33 |
| PC 37:3 | 21051.24 | 32603.56 | 22853.34 | 35487.78 |
| PC 37:4 | 301471.20 | 269943.16 | 322941.76 | 391448.82 |
| PC 37:6 | 148291.65 | 303987.76 | 189669.76 | 243554.88 |
| PC 37:7 | 2029713.54 | 2304539.91 | 2144008.57 | 3834832.16 |
| PC 38:1 | 76798.05 | 149720.78 | 98303.55 | 188080.29 |
| PC 38:10 | 6431.71 | 35423.17 | 4503.30 | 24550.39 |
| PC 38:2 | 302157.43 | 572709.16 | 331610.20 | 651275.79 |
| PC 38:3 | 1760194.47 | 1248221.68 | 1054678.72 | 1976833.20 |
| PC 38:4 | 4150984.83 | 4886451.69 | 5394808.40 | 8980959.19 |
| PC 38:5 | 1241094.77 | 2501918.94 | 2182573.00 | 1745327.31 |
| PC 38:6 | 12593848.39 | 9875153.22 | 12001374.40 | 13489484.56 |
| PC 38:7 | 6116539.11 | 7762998.33 | 7773498.53 | 9457634.20 |
| PC 38:8 | 179028.34 | 253173.75 | 187217.90 | 190748.73 |
| PC 38:9 | 43365.68 | 76444.66 | 52088.53 | 92926.17 |
| PC 39:3 | 32979.65 | 41961.40 | 24824.53 | 63542.46 |
| PC 39:4 | 153348.07 | 114281.99 | 124236.13 | 192755.47 |
| PC 39:5 | 52669.18 | 70922.42 | 65360.22 | 74848.29 |
| PC 39:6 | 1140236.11 | 817598.40 | 1214826.55 | 1058701.40 |
| PC 39:7 | 412971.43 | 313238.07 | 378118.48 | 343554.40 |
| PC 39:8 | 54039.05 | 64335.76 | 82932.68 | 124577.45 |
| PC 40:0 | 19068.09 | 26641.43 | 13106.40 | 46143.67 |
| PC 40:1 | 44282.37 | 101267.80 | 81979.77 | 187254.70 |
| PC 40:10 | 47506.90 | 56251.89 | 47453.22 | 53173.54 |
| PC 40:2 | 21255.90 | 172682.20 | 30005.52 | 229361.29 |
| PC 40:3 | 30352.82 | 55448.17 | 43752.44 | 76231.83 |
| PC 40:4 | 160763.39 | 452573.55 | 274126.10 | 650650.58 |
| PC 40:5 | 2059449.42 | 2810341.98 | 2633374.56 | 4064423.39 |
| PC 40:6 | 11675779.92 | 10221779.40 | 11189374.38 | 14594208.21 |
| PC 40:7 | 1844669.92 | 2735586.71 | 2386291.68 | 3160595.59 |
| PC 40:8 | 10252018.16 | 8769797.94 | 9672895.23 | 11296480.04 |
| PC 40:9 | 971034.14 | 961067.31 | 1185586.69 | 999682.56 |
| PC 41:6 | 307786.51 | 363130.43 | 393665.86 | 430072.36 |
| PC 41:7 | 25479.66 | 27289.41 | 29505.89 | 31911.93 |
| PC 42:1 | 97959.39 | 203821.94 | 181355.90 | 416533.56 |
| PC 42:10 | 540348.64 | 517988.86 | 784295.23 | 757646.52 |
| PC 42:11 | 146607.14 | 150657.56 | 141860.31 | 130423.03 |
| PC 42:2 | 43061.17 | 94697.93 | 66838.57 | 153338.66 |
| PC 42:4 | 7510.17 | 64586.03 | 7964.78 | 123775.60 |
| PC 42:5 | 33680.52 | 214268.73 | 39424.22 | 313359.44 |
| PC 42:6 | 145235.97 | 192623.97 | 211514.05 | 302759.17 |
| PC 42:7 | 25388.10 | 44114.59 | 32669.84 | 50884.90 |
| PC 42:8 | 18863.10 | 44862.81 | 22119.39 | 38460.47 |
| PC 42:9 | 130498.04 | 131635.84 | 174054.33 | 176928.13 |
| PC 44:10 | 19083.72 | 79270.90 | 22882.69 | 194929.72 |
| PC 44:11 | 331201.71 | 317673.38 | 417546.65 | 582437.96 |
| PC 44:12 | 273983.32 | 302322.78 | 307562.36 | 344643.49 |
| PC 44:2 | 7256.79 | 66940.92 | 7180.96 | 49428.28 |
| PC 44:4 | 9309.40 | 53324.44 | 6964.42 | 87404.53 |
| PC 44:9 | 4094.28 | 6624.38 | 7736.45 | 16410.19 |
| PE 30:0 | 36620.32 | 58535.36 | 13395.20 | 28183.76 |
| PE 30:1 | 3454.21 | 14048.44 | 673.86 | 3791.63 |
| PE 32:0 | 40428.76 | 91603.00 | 53361.59 | 79863.96 |

| | | | | |
|---|---|---|---|---|
| PE 32:1 | 15877.11 | 15661.28 | 9315.90 | 20334.76 |
| PE 32:2 | 33132.12 | 57330.04 | 9788.40 | 23706.30 |
| PE 32:3 | 10914.35 | 15488.47 | 3422.22 | 7983.80 |
| PE 33:0 | 15169.13 | 18845.84 | 12846.38 | 26888.13 |
| PE 33:1 | 5555.44 | 6295.62 | 4461.86 | 6685.02 |
| PE 33:2 | 31751.84 | 37639.15 | 17853.63 | 23764.70 |
| PE 34:1 | 81111.54 | 150618.08 | 91818.23 | 245554.68 |
| PE 34:2 | 855487.90 | 911289.36 | 548733.09 | 1181651.06 |
| PE 34:3 | 132087.02 | 184821.16 | 122978.11 | 270733.20 |
| PE 34:4 | 31331.89 | 41703.87 | 16491.59 | 21764.73 |
| PE 35:0 | 31416.08 | 32559.22 | 29987.94 | 43671.35 |
| PE 35:1 | 46252.28 | 78211.09 | 48211.23 | 99150.96 |
| PE 35:2 | 74576.55 | 90529.84 | 69299.70 | 113958.98 |
| PE 35:3 | 21190.12 | 43441.85 | 18023.52 | 32998.46 |
| PE 35:4 | 11803.81 | 18575.90 | 10672.33 | 23015.21 |
| PE 36:0 | 79566.29 | 99048.41 | 69683.49 | 117250.25 |
| PE 36:1 | 414929.18 | 1222631.19 | 592474.29 | 1678937.49 |
| PE 36:2 | 786192.53 | 1113993.76 | 766377.28 | 1485100.83 |
| PE 36:3 | 652050.98 | 1082976.17 | 657217.46 | 1642363.90 |
| PE 36:4 | 204787.73 | 186429.76 | 236425.19 | 378548.92 |
| PE 36:5 | 69989.21 | 100653.69 | 64695.64 | 143348.07 |
| PE 36:6 | 83277.29 | 96473.35 | 9776.51 | 22112.16 |
| PE 37:1 | 62276.87 | 98004.79 | 53488.44 | 123263.00 |
| PE 37:2 | 102010.78 | 115475.05 | 105296.92 | 150839.69 |
| PE 37:3 | 23476.55 | 25771.20 | 26964.33 | 45951.94 |
| PE 37:4 | 26462.64 | 28272.27 | 28856.09 | 52635.35 |
| PE 37:6 | 115315.43 | 106808.65 | 64933.40 | 64018.32 |
| PE 38:1 | 89346.95 | 127303.89 | 69011.37 | 168654.94 |
| PE 38:2 | 174436.20 | 374999.13 | 220160.46 | 479666.73 |
| PE 38:3 | 677373.43 | 1186023.49 | 808884.79 | 1580093.38 |
| PE 38:4 | 1056591.72 | 1375966.34 | 1373369.32 | 2422646.82 |
| PE 38:5 | 2832727.72 | 2869732.57 | 2902864.19 | 3384681.76 |
| PE 38:6 | 3390828.24 | 3234280.16 | 3643850.31 | 3379843.67 |
| PE 38:7 | 385442.85 | 475872.89 | 385981.82 | 768519.92 |
| PE 38:8 | 46829.42 | 64029.63 | 31550.62 | 40400.35 |
| PE 39:4 | 167158.49 | 146813.48 | 177562.68 | 271524.12 |
| PE 39:5 | 145047.37 | 177984.28 | 192741.80 | 221128.31 |
| PE 39:6 | 221208.48 | 756116.14 | 214849.43 | 796000.08 |
| PE 39:7 | 559017.77 | 516126.40 | 389122.37 | 650259.09 |
| PE 39:8 | 58113.48 | 55791.98 | 50759.81 | 50818.71 |
| PE 40:1 | 30306.86 | 58509.86 | 27140.73 | 92717.25 |
| PE 40:10 | 22895.21 | 33322.63 | 18205.33 | 36055.47 |
| PE 40:2 | 39931.00 | 73394.03 | 48895.97 | 88574.41 |
| PE 40:3 | 94588.59 | 135797.52 | 92052.19 | 157503.12 |
| PE 40:4 | 343317.21 | 925858.98 | 401219.41 | 1187877.28 |
| PE 40:5 | 7181334.24 | 4265661.06 | 5918992.22 | 7224144.89 |
| PE 40:6 | 14533180.15 | 5606262.52 | 7817142.10 | 8658803.51 |
| PE 40:7 | 2029713.54 | 2304539.91 | 2144008.57 | 3834832.16 |
| PE 40:8 | 5685145.22 | 7015603.26 | 5864173.71 | 6608702.61 |
| PE 40:9 | 421668.12 | 443543.71 | 498546.01 | 564427.46 |
| PE 41:1 | 11627.06 | 33122.71 | 11649.45 | 29502.09 |
| PE 41:5 | 178584.75 | 181828.63 | 229106.61 | 223275.71 |
| PE 41:6 | 755493.51 | 699735.62 | 886808.30 | 881936.71 |
| PE 41:7 | 85910.56 | 108553.55 | 116099.57 | 106753.18 |
| PE 42:1 | 21690.84 | 42420.29 | 15999.46 | 81772.22 |
| PE 42:10 | 128786.90 | 109308.86 | 189853.46 | 202833.13 |

| | | | | |
|---|---|---|---|---|
| PE 42:11 | 80078.00 | 92978.31 | 99522.29 | 90699.03 |
| PE 42:2 | 18214.21 | 56886.88 | 26015.93 | 80461.57 |
| PE 42:3 | 25448.54 | 49840.50 | 26220.48 | 51699.17 |
| PE 42:6 | 72338.52 | 99209.94 | 109209.36 | 119973.23 |
| PE 42:7 | 83103.06 | 83026.85 | 140286.30 | 138290.37 |
| PE 42:8 | 146294.96 | 136219.04 | 184880.24 | 223826.43 |
| PE 42:9 | 693857.20 | 889022.37 | 1016331.06 | 1039512.11 |
| PE 44:10 | 268344.62 | 329555.32 | 568325.84 | 611166.58 |
| PE 44:11 | 651585.27 | 491386.50 | 754364.12 | 946275.04 |
| PE 44:12 | 512401.68 | 482699.04 | 519644.12 | 679001.53 |
| PE 44:4 | 15541.11 | 30726.23 | 20234.85 | 43668.82 |
| PE 44:9 | 30182.76 | 26831.71 | 31717.00 | 49142.57 |
| PG 30:0 | 28130.66 | 16615.00 | 9867.73 | 15076.00 |
| PG 31:0 | 7269.69 | 7020.74 | 7780.35 | 8298.93 |
| PG 32:0 | 183655.83 | 168796.34 | 213898.35 | 213993.09 |
| PG 32:1 | 71741.63 | 72752.72 | 61042.82 | 66381.51 |
| PG 32:2 | 11612.88 | 19603.54 | 5943.38 | 11570.60 |
| PG 33:0 | 38522.96 | 44604.39 | 45808.88 | 46496.74 |
| PG 33:1 | 39405.21 | 36046.27 | 36265.44 | 27416.43 |
| PG 34:1 | 2884723.80 | 1654712.86 | 2339466.73 | 2670641.59 |
| PG 34:2 | 572272.65 | 755607.53 | 612309.69 | 628086.70 |
| PG 34:3 | 50531.29 | 34097.43 | 52155.79 | 46637.26 |
| PG 36:0 | 67671.77 | 78276.64 | 93247.15 | 113238.10 |
| PG 36:1 | 174715.70 | 166710.10 | 88948.74 | 202551.14 |
| PG 36:2 | 702961.36 | 1069817.31 | 705240.30 | 1062547.57 |
| PG 36:3 | 193064.72 | 283715.84 | 218784.43 | 291464.63 |
| PG 36:4 | 29563.41 | 62052.66 | 57546.57 | 55219.95 |
| PG 36:5 | 13527.70 | 12007.41 | 14262.78 | 24925.35 |
| PG 38:2 | 21366.27 | 53079.55 | 23698.04 | 42311.81 |
| PG 38:3 | 43226.69 | 67775.90 | 50018.64 | 92770.51 |
| PG 38:4 | 21004.32 | 43652.98 | 27060.70 | 62819.72 |
| PG 38:5 | 63707.78 | 119974.58 | 76584.48 | 106962.99 |
| PG 38:6 | 103755.31 | 144887.10 | 98880.65 | 163949.64 |
| PG 38:7 | 9949.30 | 23536.02 | 8185.08 | 19535.91 |
| PG 40:6 | 91303.34 | 101579.33 | 92787.49 | 95578.39 |
| PG 40:7 | 83738.15 | 181171.91 | 93270.97 | 196991.93 |
| PG 40:8 | 57953.57 | 167459.15 | 79688.39 | 142434.97 |
| PG 40:9 | 6521.03 | 9523.46 | 3505.87 | 13063.71 |
| PG 42:10 | 6482.01 | 28172.95 | 11486.25 | 42405.93 |
| PG 42:6 | 878775.00 | 825180.61 | 971516.59 | 731121.96 |
| PG 42:8 | 13163.86 | 41174.64 | 18348.44 | 43902.77 |
| PG 42:9 | 13639.62 | 31407.38 | 13969.71 | 30400.23 |
| PG 44:10 | 10422.75 | 30712.51 | 9606.50 | 37609.96 |
| PG 44:11 | 21393.43 | 77517.88 | 31390.51 | 70854.71 |
| PG 44:12 | 58899.44 | 126728.63 | 43808.56 | 117055.25 |
| PG 44:9 | 131623.96 | 64013.71 | 96898.03 | 161248.37 |
| PI 34:2 | 90862.09 | 171883.79 | 90034.56 | 165456.84 |
| PI 36:2 | 193665.63 | 404600.17 | 225864.52 | 549620.86 |
| PI 36:3 | 67121.21 | 107243.70 | 66634.32 | 158201.49 |
| PI 36:4 | 55316.98 | 97135.64 | 66264.01 | 135937.25 |
| PI 37:4 | 194188.81 | 156246.19 | 174090.65 | 237620.09 |
| PI 38:3 | 899798.39 | 1010184.13 | 918455.51 | 1754460.47 |
| PI 38:4 | 2678640.90 | 3270924.45 | 3704684.76 | 4403254.82 |
| PI 38:5 | 186208.91 | 283084.87 | 346724.07 | 423309.12 |
| PI 39:4 | 46780.62 | 46879.69 | 53288.30 | 81531.50 |
| PI 40:5 | 393887.89 | 373353.64 | 475866.95 | 678051.42 |

| | | | | |
|---|---|---|---|---|
| PI 40:6 | 986325.72 | 817430.12 | 1254750.23 | 1432739.36 |
| PI 40:7 | 67956.73 | 56493.98 | 66809.97 | 116070.33 |
| Plasmenyl-PC 24:0 | 1265.79 | 3709.01 | 2167.19 | 5315.91 |
| Plasmenyl-PC 32:0 | 157040.19 | 175552.73 | 149519.90 | 190430.07 |
| Plasmenyl-PC 33:0 | 7243.32 | 12662.43 | 6039.04 | 19770.56 |
| Plasmenyl-PC 33:5 | 19480.95 | 28411.16 | 17348.09 | 35227.18 |
| Plasmenyl-PC 34:0 | 53658.58 | 88682.96 | 70931.49 | 87596.70 |
| Plasmenyl-PC 34:1 | 74745.53 | 140438.61 | 63222.12 | 188794.21 |
| Plasmenyl-PC 34:2 | 1819.40 | 4950.28 | 2205.50 | 6431.92 |
| Plasmenyl-PC 35:1 | 11551.04 | 24397.82 | 15725.84 | 22962.66 |
| Plasmenyl-PC 35:3 | 6442.15 | 4603.93 | 4859.22 | 7845.89 |
| Plasmenyl-PC 36:0 | 24741.52 | 39512.22 | 24522.95 | 31056.47 |
| Plasmenyl-PC 36:1 | 33264.48 | 222424.30 | 35224.54 | 259375.59 |
| Plasmenyl-PC 36:2 | 7937.79 | 11528.97 | 8996.36 | 13112.76 |
| Plasmenyl-PC 36:3 | 55283.41 | 85944.15 | 45686.84 | 148997.83 |
| Plasmenyl-PC 36:4 | 32682.80 | 60692.50 | 37573.49 | 67182.35 |
| Plasmenyl-PC 36:5 | 51717.62 | 68488.09 | 39538.70 | 93928.87 |
| Plasmenyl-PC 36:6 | 104543.57 | 99042.95 | 127440.23 | 129540.35 |
| Plasmenyl-PC 37:2 | 66534.36 | 74228.40 | 60405.21 | 93925.86 |
| Plasmenyl-PC 37:3 | 7317.59 | 9606.50 | 7366.35 | 13022.63 |
| Plasmenyl-PC 37:4 | 8493.17 | 17462.22 | 7524.38 | 27780.34 |
| Plasmenyl-PC 38:0 | 8882.47 | 11000.86 | 7695.44 | 12641.14 |
| Plasmenyl-PC 38:3 | 46737.84 | 148976.81 | 43005.78 | 234832.23 |
| Plasmenyl-PC 38:4 | 135245.81 | 205736.71 | 97367.62 | 308103.55 |
| Plasmenyl-PC 38:5 | 575082.93 | 661958.62 | 634557.29 | 866046.52 |
| Plasmenyl-PC 38:6 | 423756.92 | 576297.80 | 412685.07 | 514194.40 |
| Plasmenyl-PC 39:3 | 59201.55 | 70609.63 | 58281.65 | 93667.23 |
| Plasmenyl-PC 39:5 | 36637.86 | 33732.83 | 34725.95 | 44353.83 |
| Plasmenyl-PC 39:6 | 36606.22 | 35224.47 | 39634.03 | 47974.43 |
| Plasmenyl-PC 40:0 | 4109.40 | 24759.02 | 3685.48 | 27010.34 |
| Plasmenyl-PC 40:1 | 3240.13 | 28155.90 | 3515.01 | 18725.23 |
| Plasmenyl-PC 40:2 | 4749.56 | 11139.37 | 8482.98 | 11574.50 |
| Plasmenyl-PC 40:3 | 29481.28 | 32396.50 | 36692.47 | 57069.91 |
| Plasmenyl-PC 40:4 | 1321.01 | 17161.53 | 1306.26 | 12870.42 |
| Plasmenyl-PC 40:5 | 98747.62 | 168540.86 | 111842.02 | 214667.43 |
| Plasmenyl-PC 40:6 | 93315.37 | 177277.77 | 145418.18 | 206664.32 |
| Plasmenyl-PC 41:3 | 5918.28 | 36767.94 | 3683.62 | 31997.71 |
| Plasmenyl-PC 42:1 | 4210.34 | 12263.82 | 3834.31 | 7501.56 |
| Plasmenyl-PC 42:3 | 34936.54 | 48479.80 | 37802.02 | 51867.35 |
| Plasmenyl-PC 42:4 | 5889.22 | 8395.59 | 7206.32 | 19105.27 |
| Plasmenyl-PC 42:5 | 21211.90 | 20974.86 | 17963.53 | 37203.21 |
| Plasmenyl-PC 42:6 | 14388.09 | 13890.49 | 4243.31 | 15075.21 |
| Plasmenyl-PC 44:3 | 3583.27 | 9824.06 | 2049.62 | 10271.34 |
| Plasmenyl-PC 44:4 | 5026.79 | 34140.30 | 4875.61 | 47952.99 |
| Plasmenyl-PC 44:5 | 4472.80 | 10295.89 | 3914.21 | 11022.64 |
| Plasmenyl-PC 44:6 | 14289.04 | 19863.01 | 14927.14 | 37724.36 |
| Plasmenyl-PC 46:4 | 2430.30 | 12837.97 | 702.73 | 13121.48 |
| Plasmenyl-PE 30:0 | 6359.45 | 10948.58 | 2676.99 | 4387.67 |
| Plasmenyl-PE 32:0 | 144245.47 | 183355.25 | 113327.00 | 212629.77 |
| Plasmenyl-PE 32:1 | 33413.47 | 78664.80 | 23992.22 | 56623.51 |
| Plasmenyl-PE 32:2 | 22267.34 | 42149.43 | 5003.15 | 8784.38 |
| Plasmenyl-PE 33:0 | 14159.38 | 24374.34 | 14934.13 | 20647.35 |
| Plasmenyl-PE 33:1 | 10623.02 | 15111.70 | 7503.69 | 13834.87 |
| Plasmenyl-PE 34:0 | 702284.18 | 925895.92 | 724954.66 | 1016039.82 |
| Plasmenyl-PE 34:1 | 2010029.55 | 1412770.46 | 1698358.82 | 2655432.43 |
| Plasmenyl-PE 34:2 | 109434.34 | 185954.93 | 62888.85 | 242653.35 |

| | | | | |
|---|---|---|---|---|
| Plasmenyl-PE 34:3 | 16054.76 | 32713.71 | 9695.28 | 28538.23 |
| Plasmenyl-PE 34:4 | 14969.95 | 20584.28 | 8833.47 | 29665.22 |
| Plasmenyl-PE 35:1 | 86577.30 | 203354.94 | 142441.68 | 257614.21 |
| Plasmenyl-PE 35:2 | 34614.13 | 47129.87 | 29763.74 | 50985.66 |
| Plasmenyl-PE 35:4 | 4033.29 | 11899.87 | 4957.83 | 11007.87 |
| Plasmenyl-PE 36:0 | 111531.94 | 137126.26 | 113614.37 | 154425.72 |
| Plasmenyl-PE 36:1 | 2904842.64 | 3810921.06 | 3573599.35 | 4893221.08 |
| Plasmenyl-PE 36:2 | 430114.95 | 365920.61 | 283605.92 | 565429.12 |
| Plasmenyl-PE 36:3 | 258699.01 | 507688.04 | 262977.47 | 633726.69 |
| Plasmenyl-PE 36:4 | 1320852.22 | 1651377.43 | 1253915.38 | 2905432.17 |
| Plasmenyl-PE 36:5 | 92237.32 | 210442.43 | 94533.11 | 194797.02 |
| Plasmenyl-PE 36:6 | 48917.61 | 54117.89 | 27001.53 | 46968.25 |
| Plasmenyl-PE 37:0 | 12731.11 | 25754.47 | 17169.21 | 44776.88 |
| Plasmenyl-PE 37:1 | 90161.96 | 65223.52 | 65057.03 | 119387.23 |
| Plasmenyl-PE 37:2 | 28714.14 | 50892.11 | 30597.42 | 46146.01 |
| Plasmenyl-PE 37:4 | 50065.72 | 106186.71 | 56819.56 | 135678.92 |
| Plasmenyl-PE 37:5 | 26481.25 | 49946.21 | 26125.27 | 37462.54 |
| Plasmenyl-PE 37:6 | 73691.44 | 84152.48 | 62153.09 | 84482.70 |
| Plasmenyl-PE 38:1 | 159168.27 | 252318.13 | 171322.38 | 292966.79 |
| Plasmenyl-PE 38:2 | 95575.62 | 227710.65 | 119720.93 | 230851.52 |
| Plasmenyl-PE 38:3 | 143716.34 | 451034.17 | 235215.77 | 580621.14 |
| Plasmenyl-PE 38:4 | 1275039.98 | 1521995.09 | 1264689.44 | 2614620.36 |
| Plasmenyl-PE 38:5 | 626766.80 | 745738.04 | 594616.95 | 966553.15 |
| Plasmenyl-PE 38:6 | 13649404.87 | 16405762.66 | 13348674.61 | 16330193.34 |
| Plasmenyl-PE 39:4 | 49352.39 | 90288.84 | 48570.25 | 116037.13 |
| Plasmenyl-PE 39:5 | 210106.16 | 225606.81 | 198058.05 | 256897.31 |
| Plasmenyl-PE 39:6 | 441813.48 | 507205.37 | 455444.90 | 558504.48 |
| Plasmenyl-PE 40:2 | 33065.85 | 62651.88 | 29356.28 | 88493.34 |
| Plasmenyl-PE 40:3 | 134107.99 | 178658.74 | 155066.25 | 298379.85 |
| Plasmenyl-PE 40:4 | 553610.44 | 1431889.43 | 836735.59 | 1588871.28 |
| Plasmenyl-PE 40:5 | 494238.67 | 702273.32 | 560989.00 | 767859.90 |
| Plasmenyl-PE 40:6 | 10792312.64 | 5384178.01 | 8168344.09 | 8033016.17 |
| Plasmenyl-PE 41:6 | 147284.47 | 253025.10 | 194656.85 | 226321.51 |
| Plasmenyl-PE 42:4 | 73324.87 | 98728.02 | 71219.40 | 160258.07 |
| Plasmenyl-PE 42:5 | 221207.69 | 268184.89 | 232318.07 | 355568.29 |
| Plasmenyl-PE 42:6 | 1215138.86 | 852785.69 | 890815.40 | 1220419.17 |
| PS 34:0 | 2820540.23 | 2205198.39 | 2469263.76 | 3308217.87 |
| PS 36:0 | 4139824.94 | 3092002.24 | 3901391.48 | 4818180.22 |
| PS 36:1 | 47142.82 | 99178.02 | 48593.14 | 129250.36 |
| PS 38:3 | 36401.39 | 50329.89 | 42585.22 | 98604.00 |
| PS 38:4 | 77491.62 | 97030.45 | 91639.83 | 234270.28 |
| PS 38:6 | 139885.69 | 126713.85 | 190048.13 | 131174.66 |
| PS 39:6 | 54307.88 | 45212.04 | 62060.97 | 54568.51 |
| PS 40:5 | 879962.62 | 1801312.98 | 1196654.22 | 2246878.09 |
| PS 40:6 | 1473338.50 | 1132195.93 | 1218207.83 | 1567315.27 |
| PS 40:7 | 51331.62 | 84697.08 | 63491.73 | 102428.66 |
| PS 40:8 | 24798.47 | 50654.99 | 36303.51 | 52582.11 |
| PS 40:9 | 4326.60 | 6359.42 | 5440.68 | 8345.94 |
| PS 41:6 | 243166.35 | 410967.50 | 360270.77 | 463578.31 |
| PS 42:1 | 114235.24 | 199367.59 | 141370.67 | 271054.87 |
| PS 42:10 | 19790.42 | 32699.11 | 24745.91 | 39266.95 |
| PS 42:11 | 57970.39 | 51438.41 | 53923.23 | 62458.83 |
| PS 42:2 | 5623.92 | 8580.07 | 7667.41 | 10549.06 |
| PS 42:7 | 60996.81 | 165488.07 | 60424.99 | 140739.08 |
| PS 44:10 | 32331.76 | 41380.76 | 55867.49 | 81515.31 |
| PS 44:11 | 137451.35 | 134414.15 | 170762.15 | 175183.55 |

| | | | | |
|---|---|---|---|---|
| PS 44:12 | 20400.87 | 21094.94 | 30682.82 | 32492.78 |
| SM 30:1 | 2366.01 | 4217.83 | 2931.32 | 3562.77 |
| SM 30:2 | 295.51 | 3533.93 | 630.22 | 4392.75 |
| SM 31:1 | 1877.47 | 2162.99 | 2624.98 | 2383.47 |
| SM 32:0 | 3666.14 | 2852.90 | 1952.55 | 3823.99 |
| SM 32:1 | 7487.75 | 77737.31 | 9843.90 | 90275.23 |
| SM 32:2 | 1294.16 | 1202.01 | 1673.30 | 5591.97 |
| SM 33:1 | 14166.80 | 35800.91 | 21291.56 | 59849.67 |
| SM 33:2 | 3033.17 | 2735.79 | 2993.95 | 4565.21 |
| SM 34:0 | 31074.70 | 59450.26 | 36137.29 | 67553.89 |
| SM 34:1 | 1888318.41 | 3733473.68 | 1691956.22 | 6111029.63 |
| SM 34:2 | 64913.40 | 273575.43 | 113410.52 | 216333.84 |
| SM 34:3 | 943.35 | 2055.26 | 1670.13 | 4243.51 |
| SM 35:2 | 4587.12 | 11570.25 | 5158.92 | 9739.85 |
| SM 36:1 | 3398530.47 | 2935446.74 | 3997036.77 | 4598054.56 |
| SM 36:2 | 43971.51 | 219329.28 | 57733.75 | 403224.74 |
| SM 36:3 | 3420.07 | 22816.76 | 5878.17 | 33901.11 |
| SM 36:4 | 2034.73 | 1589.26 | 1090.23 | 2626.10 |
| SM 37:1 | 210567.35 | 66387.15 | 148262.72 | 123213.62 |
| SM 38:0 | 37464.68 | 68539.57 | 59893.45 | 124742.82 |
| SM 38:1 | 230622.89 | 385370.81 | 377258.42 | 694038.82 |
| SM 38:2 | 25803.38 | 43388.79 | 22642.43 | 99078.58 |
| SM 38:3 | 2145.68 | 1484.02 | 583.06 | 1499.92 |
| SM 39:1 | 58294.40 | 110193.32 | 74209.50 | 130340.27 |
| SM 40:1 | 411924.09 | 702088.78 | 505526.46 | 967536.12 |
| SM 40:2 | 39519.57 | 39866.33 | 51264.03 | 87412.10 |
| SM 40:3 | 2348.91 | 2987.17 | 1742.50 | 5086.34 |
| SM 41:1 | 377247.11 | 686392.37 | 441244.56 | 779097.39 |
| SM 41:2 | 43149.28 | 427212.85 | 49415.49 | 442560.34 |
| SM 41:5 | 15002.47 | 16752.89 | 13716.87 | 20744.41 |
| SM 41:6 | 5110.87 | 5014.49 | 2931.36 | 6329.15 |
| SM 42:1 | 464432.10 | 1115594.86 | 528646.47 | 1225962.32 |
| SM 42:2 | 701348.21 | 1988867.33 | 759655.14 | 2120053.36 |
| SM 42:3 | 391332.64 | 963969.09 | 381229.47 | 1017318.84 |
| SM 42:4 | 23861.29 | 59340.85 | 15954.71 | 63488.20 |
| SM 42:5 | 6312.74 | 7674.34 | 3378.74 | 7264.51 |
| SM 43:1 | 30163.22 | 64576.14 | 31956.79 | 72134.20 |
| SM 43:2 | 14254.20 | 30902.17 | 15911.92 | 30964.45 |
| SM 43:3 | 7358.16 | 6419.63 | 2838.07 | 6204.59 |
| SM 44:1 | 8112.30 | 65211.55 | 8879.02 | 81830.85 |
| SM 44:2 | 6945.76 | 23424.37 | 10715.69 | 22393.33 |
| SM 44:4 | 310273.05 | 520666.80 | 394121.18 | 484871.08 |
| SM 44:5 | 12141.67 | 19924.27 | 7560.06 | 12381.23 |
| TG 36:0 | 63535.31 | 400111.11 | 83574.52 | 298985.69 |
| TG 38:0 | 43930.97 | 154423.15 | 39132.49 | 220909.00 |
| TG 40:0 | 19388.46 | 43175.15 | 21687.39 | 52715.41 |
| TG 40:1 | 123113.47 | 266903.75 | 80857.10 | 341441.51 |
| TG 42:0 | 7166.42 | 8107.58 | 5942.76 | 11146.60 |
| TG 42:1 | 68845.30 | 292798.66 | 39979.56 | 367718.17 |
| TG 42:2 | 62188.70 | 285454.64 | 55019.06 | 322019.97 |
| TG 42:3 | 31140.17 | 599765.23 | 32815.07 | 697053.40 |
| TG 43:0 | 3356.55 | 30985.12 | 3398.18 | 26887.56 |
| TG 44:0 | 43423.75 | 260184.54 | 31871.97 | 386747.41 |
| TG 44:1 | 109188.55 | 314548.34 | 89418.69 | 384168.40 |
| TG 44:2 | 66497.82 | 302460.40 | 56791.94 | 344702.81 |
| TG 44:3 | 43582.16 | 162213.54 | 34036.49 | 190400.76 |

| | | | | |
|---|---|---|---|---|
| TG 45:0 | 12628.01 | 8107.05 | 7159.41 | 12874.75 |
| TG 45:1 | 12040.63 | 94682.52 | 11744.79 | 112548.01 |
| TG 45:2 | 11486.68 | 180594.54 | 10307.14 | 127937.60 |
| TG 46:0 | 159706.75 | 204051.30 | 120442.08 | 270995.48 |
| TG 46:1 | 472886.02 | 644363.81 | 362880.50 | 1097158.93 |
| TG 46:2 | 547914.44 | 750739.32 | 302552.82 | 1079006.98 |
| TG 46:3 | 121971.91 | 484095.11 | 104968.27 | 537653.12 |
| TG 46:4 | 21028.81 | 460828.88 | 18641.91 | 467218.89 |
| TG 47:0 | 14710.85 | 16313.29 | 15413.20 | 23671.09 |
| TG 47:1 | 29988.54 | 169389.03 | 29486.42 | 238433.37 |
| TG 47:2 | 43709.98 | 102737.41 | 39539.96 | 126433.61 |
| TG 47:3 | 29235.85 | 92445.29 | 17829.29 | 70369.85 |
| TG 47:4 | 10648.16 | 44406.74 | 7883.49 | 28826.65 |
| TG 48:0 | 378280.50 | 574325.41 | 292750.03 | 651752.48 |
| TG 48:1 | 1993058.20 | 1504332.25 | 1121313.59 | 2327951.64 |
| TG 48:2 | 1710804.03 | 2908704.48 | 1500209.17 | 4102149.85 |
| TG 48:3 | 973184.34 | 3208145.04 | 1233478.41 | 3221433.78 |
| TG 48:4 | 190434.10 | 766327.87 | 165744.78 | 834090.25 |
| TG 49:0 | 24117.43 | 30339.12 | 21085.62 | 32582.72 |
| TG 49:1 | 89137.19 | 140338.91 | 78863.91 | 202329.99 |
| TG 49:2 | 188180.53 | 153483.19 | 111785.88 | 261429.86 |
| TG 49:3 | 85920.38 | 198400.36 | 80250.05 | 232903.54 |
| TG 49:4 | 48798.09 | 163922.66 | 31512.98 | 149467.99 |
| TG 49:5 | 12835.70 | 72885.03 | 6664.25 | 32752.96 |
| TG 50:0 | 358192.97 | 641468.43 | 313463.15 | 774979.34 |
| TG 50:1 | 3148533.23 | 5251570.03 | 2637616.05 | 6840265.66 |
| TG 50:2 | 4769922.95 | 7711237.94 | 4014366.61 | 10508341.13 |
| TG 50:3 | 3531109.08 | 6424633.70 | 3206383.05 | 8821842.58 |
| TG 50:4 | 1525012.71 | 3764209.17 | 1468934.74 | 4706473.91 |
| TG 50:5 | 283917.79 | 1122696.03 | 260881.43 | 1287130.87 |
| TG 50:6 | 64546.21 | 155804.73 | 37211.37 | 201543.89 |
| TG 51:0 | 804278.69 | 647693.36 | 719505.74 | 992211.96 |
| TG 51:1 | 110938.66 | 228229.10 | 88314.82 | 275074.39 |
| TG 51:2 | 197297.84 | 361911.42 | 167241.12 | 467541.07 |
| TG 51:3 | 3419.77 | 15523.05 | 3163.72 | 16644.92 |
| TG 51:4 | 121173.08 | 259031.41 | 114428.04 | 352460.34 |
| TG 51:5 | 63375.91 | 129412.27 | 35000.18 | 140273.37 |
| TG 51:6 | 16940.98 | 128897.93 | 12574.67 | 65752.67 |
| TG 52:1 | 1205526.98 | 2795461.71 | 1001798.54 | 3918183.61 |
| TG 52:2 | 4451064.83 | 8629523.86 | 3565190.91 | 11011873.92 |
| TG 52:3 | 6269337.68 | 11179242.75 | 5314391.78 | 14340327.14 |
| TG 52:4 | 5143941.30 | 10078973.15 | 4357629.12 | 12165003.87 |
| TG 52:5 | 306180.07 | 1167064.51 | 304468.91 | 1641602.64 |
| TG 52:6 | 425386.09 | 1576113.22 | 388244.51 | 1834365.60 |
| TG 52:7 | 78583.49 | 317033.74 | 67560.20 | 409918.35 |
| TG 53:0 | 39556.15 | 42524.29 | 23194.00 | 63377.80 |
| TG 53:1 | 51923.08 | 149069.48 | 39262.83 | 145413.94 |
| TG 53:2 | 149386.22 | 350731.32 | 115094.34 | 361784.99 |
| TG 53:3 | 181830.89 | 364114.05 | 159212.94 | 391902.27 |
| TG 53:4 | 131607.07 | 280067.92 | 103732.82 | 320067.60 |
| TG 53:5 | 95382.80 | 135794.45 | 68564.91 | 222925.99 |
| TG 53:6 | 28919.47 | 97265.38 | 25954.98 | 134811.66 |
| TG 54:1 | 202256.72 | 617021.12 | 173313.28 | 954966.53 |
| TG 54:2 | 1045296.23 | 2787444.74 | 794383.57 | 3365190.63 |
| TG 54:3 | 2623706.35 | 5706012.91 | 2028377.81 | 6175958.32 |
| TG 54:4 | 3228853.58 | 6346149.17 | 2739301.00 | 7053559.40 |

| | | | | |
|---|---|---|---|---|
| TG 54:5 | 2959535.21 | 5847196.05 | 2477766.42 | 6468024.60 |
| TG 54:6 | 1721837.81 | 4509710.99 | 1344371.02 | 4686981.45 |
| TG 54:7 | 632889.03 | 1889533.65 | 534678.62 | 2599654.02 |
| TG 54:8 | 131793.84 | 551735.97 | 121418.38 | 694063.04 |
| TG 55:2 | 41232.40 | 157351.40 | 28251.03 | 127346.12 |
| TG 55:3 | 87751.40 | 180057.32 | 59896.29 | 186584.30 |
| TG 55:4 | 74076.86 | 98127.79 | 41022.25 | 102310.53 |
| TG 55:5 | 18441.80 | 134420.53 | 18285.92 | 159932.62 |
| TG 56:1 | 32584.45 | 111006.86 | 17769.01 | 282763.97 |
| TG 56:2 | 113868.24 | 348080.21 | 81816.98 | 420136.87 |
| TG 56:3 | 273368.02 | 713564.46 | 198152.27 | 792808.80 |
| TG 56:4 | 247132.09 | 2003547.26 | 212036.45 | 2454424.06 |
| TG 56:5 | 242663.55 | 482838.52 | 188118.53 | 632173.43 |
| TG 56:6 | 276647.32 | 512233.42 | 232580.74 | 799805.57 |
| TG 56:7 | 609609.98 | 627685.64 | 309545.49 | 1145724.42 |
| TG 56:8 | 452060.00 | 1417032.32 | 480878.12 | 1580259.33 |
| TG 56:9 | 177404.75 | 327468.43 | 106252.88 | 547324.83 |
| TG 57:3 | 9945.23 | 54125.83 | 5402.14 | 44523.49 |
| TG 57:4 | 11060.19 | 33189.43 | 6284.57 | 29823.40 |
| TG 58:1 | 6143.97 | 16212.99 | 3510.69 | 24692.11 |
| TG 58:10 | 97889.50 | 285923.35 | 81802.51 | 409823.13 |
| TG 58:11 | 18594.99 | 323697.76 | 16437.60 | 463948.58 |
| TG 58:3 | 42664.98 | 119483.21 | 36197.02 | 153597.25 |
| TG 58:4 | 35948.21 | 112243.59 | 29565.07 | 126727.40 |
| TG 58:5 | 35275.94 | 59262.76 | 27963.56 | 113989.05 |
| TG 58:6 | 49738.55 | 73400.68 | 50093.13 | 136573.50 |
| TG 58:7 | 102614.46 | 155345.20 | 85930.53 | 281837.96 |
| TG 58:8 | 167356.56 | 282229.21 | 148546.79 | 458664.04 |
| TG 58:9 | 246387.19 | 269611.83 | 133935.48 | 485513.99 |
| TG 60:10 | 16230.06 | 21846.65 | 15680.16 | 46206.27 |
| TG 60:11 | 19075.54 | 38682.13 | 15133.10 | 67251.05 |
| TG 60:12 | 34373.12 | 32807.86 | 16508.70 | 62347.16 |
| TG 60:7 | 11501.58 | 61354.33 | 15357.30 | 106572.86 |
| TG 60:9 | 13605.38 | 27972.09 | 14997.11 | 40183.25 |
| TG 62:12 | 8621.08 | 8884.18 | 7579.18 | 20529.71 |
| TG 62:13 | 8149.96 | 15757.33 | 7284.98 | 30695.51 |
| TG 62:14 | 6144.82 | 11369.51 | 5102.75 | 24933.86 |
| Unknown Cer[NDS] 42:0; [ | 4184.10 | 11330.35 | 5573.09 | 11055.42 |
| Unknown Cer[NS] 35:1; [M | 42597.33 | 32909.31 | 36057.84 | 46444.72 |
| Unknown Cer[NS] 40:1; [M | 2921.16 | 4377.48 | 2713.50 | 3951.43 |
| Unknown Cer[NS] 42:1; [M | 16028.74 | 33103.55 | 15995.30 | 31888.71 |
| Unknown Cer[NS] 42:2; [M | 100441.11 | 188546.85 | 86205.29 | 193626.63 |
| Unknown LysoPC 16:0; [M | 7064.28 | 10519.78 | 11565.73 | 18524.97 |
| Unknown PA 38:6; [M-H]-@ | 30281.31 | 23062.11 | 23230.72 | 24168.43 |
| Unknown PC 34:0; [M+Had | 72104.98 | 129306.02 | 89750.89 | 109291.77 |
| Unknown PC 34:1; [M+Had | 93251.92 | 94987.94 | 109170.31 | 119898.58 |
| Unknown PC 34:2; [M+H]+ | 3975.79 | 9897.30 | 2321.92 | 13156.27 |
| Unknown PC 34:3; [M+H]+ | 7246.23 | 27007.06 | 3028.18 | 24782.39 |
| Unknown PC 36:1; [M+H]+ | 6403.01 | 7765.62 | 5189.23 | 8758.53 |
| Unknown PC 36:2; [M+H]+ | 20401.75 | 15845.02 | 10523.97 | 38000.78 |
| Unknown PC 36:3; [M+H]+ | 15194.52 | 25294.99 | 9879.44 | 47221.58 |
| Unknown PC 36:4; [M+H]+ | 7812.01 | 28929.09 | 6847.14 | 21673.64 |
| Unknown PC 36:5; [M+H]+ | 4804.54 | 18104.18 | 4207.05 | 21601.15 |
| Unknown PC 38:4; [M+H]+ | 3749.91 | 3681.99 | 2780.30 | 6528.62 |
| Unknown PC 39:8; [M+H]+ | 82571.83 | 71017.59 | 92500.24 | 68169.59 |
| Unknown PE 34:0; [M-H]-@ | 82979.90 | 69113.77 | 56724.68 | 111404.88 |

| | | | | |
|---|---|---|---|---|
| Unknown PE 34:1; [M-H]-@ | 43552.46 | 55246.12 | 40435.41 | 69972.13 |
| Unknown PE 34:2; [M-H]-@ | 30550.77 | 23638.29 | 18691.44 | 35179.89 |
| Unknown PE 38:4; [M-H]-@ | 109066.58 | 129097.56 | 114067.86 | 223045.15 |
| Unknown PE 40:6; [M-H]-@ | 569740.03 | 764336.35 | 777575.64 | 733980.67 |
| Unknown PE 40:7; [M+H]+ | 87636.73 | 97756.68 | 65793.01 | 87604.17 |
| Unknown PG 34:0; [M-H]-@ | 635742.29 | 790978.24 | 977333.23 | 1556867.21 |
| Unknown PG 34:3; [M-H]-@ | 50531.29 | 36908.27 | 52155.79 | 46637.26 |
| Unknown PG 36:2; [M-H]-@ | 702961.36 | 1069817.31 | 705240.30 | 1062547.57 |
| Unknown PG 36:4; [M-H]-@ | 59986.61 | 188058.86 | 62925.02 | 144377.04 |
| Unknown PG 36:5; [M-H]-@ | 4589.16 | 33740.46 | 5801.89 | 17240.16 |
| Unknown PG 38:4; [M-H]-@ | 84291.45 | 138988.52 | 118645.93 | 177599.10 |
| Unknown PG 38:5; [M-H]-@ | 78681.29 | 68690.14 | 69366.21 | 125570.34 |
| Unknown PG 38:6; [M-H]-@ | 18867.24 | 62261.76 | 28733.44 | 78583.15 |
| Unknown PG 40:6; [M-H]-@ | 69857.49 | 84125.41 | 57471.21 | 125036.31 |
| Unknown Plasmenyl-PC 38 | 62454.28 | 80650.11 | 78231.28 | 97091.51 |
| Unknown Plasmenyl-PC 40 | 60604.00 | 69334.65 | 84219.61 | 106065.07 |
| Unknown Plasmenyl-PE 37 | 19126.32 | 34064.63 | 19195.22 | 38415.78 |
| Unknown Plasmenyl-PE 38 | 55857.27 | 120139.45 | 65809.64 | 98944.46 |
| Unknown Plasmenyl-PE 40 | 565680.76 | 962253.43 | 903243.44 | 1158500.40 |
| Unknown PS 34:0; [M-H]-@ | 41987.93 | 33379.47 | 27268.92 | 50422.95 |
| Unknown PS 40:6; [M-H]-@ | 50897.66 | 41429.85 | 35699.57 | 48139.28 |
| Unknown SM 40:1; [M+H]+ | 499883.32 | 660781.46 | 497826.36 | 894881.59 |
| Unknown SM 42:2; [M+Ha | 622041.74 | 357822.18 | 336981.22 | 867529.14 |
| Unknown TG 48:0; [M+Na] | 11407.70 | 34897.96 | 7202.66 | 34751.95 |
| Unknown TG 50:1; [M+Na] | 16913.10 | 68823.98 | 13569.56 | 42107.83 |
| Unknown TG 51:3; [M+NH | 218729.50 | 299040.68 | 201730.71 | 442361.05 |
| Unknown Unknown_FAHF | 115597.90 | 208935.14 | 127499.91 | 213063.29 |
| Unknown_FAHFA 30:2 | 102573.76 | 86747.07 | 96160.32 | 148948.93 |
| Unknown_FAHFA 31:0 | 5068.03 | 13410.95 | 4830.26 | 18805.71 |
| Unknown_FAHFA 32:1 | 8119.57 | 42058.37 | 10037.18 | 56468.64 |
| Unknown_FAHFA 32:3 | 4920.08 | 16155.20 | 8858.94 | 27281.62 |
| Unknown_FAHFA 34:2 | 4908.75 | 11947.56 | 6953.32 | 17721.44 |
| Unknown_FAHFA 34:4 | 12121.09 | 59506.92 | 16213.06 | 92635.94 |
| Unknown_FAHFA 34:5 | 57441.85 | 54873.04 | 42989.60 | 72076.20 |
| Unknown_FAHFA 35:0 | 39484.65 | 37764.06 | 37143.96 | 59737.72 |
| Unknown_FAHFA 35:3 | 9531.42 | 7983.94 | 9687.17 | 12889.07 |
| Unknown_FAHFA 35:4 | 33942.27 | 39947.10 | 34088.23 | 38383.69 |
| Unknown_FAHFA 36:0 | 27454.53 | 27379.65 | 24311.29 | 37386.93 |
| Unknown_FAHFA 36:2 | 13710.00 | 12513.62 | 14044.74 | 20975.01 |
| Unknown_FAHFA 36:3 | 37409.35 | 11390.46 | 22435.15 | 21723.35 |
| Unknown_FAHFA 36:4 | 22373.99 | 57272.07 | 25511.58 | 63937.13 |
| Unknown_FAHFA 36:5 | 5112.14 | 12801.41 | 8092.97 | 14474.63 |
| Unknown_FAHFA 36:7 | 13882.62 | 16144.43 | 12156.67 | 28149.64 |
| Unknown_FAHFA 37:3 | 44095.02 | 82619.00 | 56690.62 | 102844.50 |
| Unknown_FAHFA 38:3 | 27961.53 | 17223.22 | 23836.78 | 35520.21 |
| Unknown_FAHFA 38:4 | 13097.82 | 18037.03 | 14672.22 | 34029.39 |
| Unknown_FAHFA 38:5 | 25973.47 | 95403.35 | 33930.93 | 97511.30 |
| Unknown_FAHFA 38:6 | 267125.24 | 575398.74 | 306252.89 | 619179.53 |
| Unknown_FAHFA 38:8 | 26952.74 | 32041.21 | 23629.65 | 52595.58 |
| Unknown_FAHFA 39:3 | 52974.89 | 43957.54 | 43841.68 | 48896.94 |
| Unknown_FAHFA 40:1 | 2565.90 | 5194.57 | 1966.98 | 3780.36 |
| Unknown_FAHFA 40:3 | 402967.58 | 188684.09 | 262392.52 | 317705.86 |
| Unknown_FAHFA 40:4 | 70301.45 | 114326.46 | 85112.02 | 133048.59 |
| Unknown_FAHFA 40:5 | 58907.66 | 75031.32 | 51439.80 | 78011.07 |
| Unknown_FAHFA 40:6 | 17923.37 | 43729.21 | 27972.16 | 56261.26 |
| Unknown_FAHFA 40:7 | 13460.00 | 35384.68 | 19784.32 | 89712.38 |

| | | | | |
|---|---|---|---|---|
| Unknown_FAHFA 40:8 | 42334.80 | 182682.78 | 76453.51 | 235963.49 |
| Unknown_FAHFA 40:9 | 24886.02 | 104421.97 | 40772.24 | 140221.62 |
| Unknown_FAHFA 41:3 | 5678.13 | 7769.45 | 6595.02 | 18133.66 |
| Unknown_FAHFA 42:2 | 3038.21 | 6065.22 | 4886.70 | 7267.29 |
| Unknown_FAHFA 42:3 | 10702.75 | 14856.15 | 9800.67 | 21727.81 |
| Unknown_FAHFA 42:4 | 15048.43 | 16102.67 | 9859.32 | 19238.07 |
| Unknown_FAHFA 42:7 | 9840.68 | 16514.34 | 12792.08 | 23224.28 |
| Unknown_FAHFA 42:9 | 24899.85 | 87201.53 | 40822.79 | 68416.16 |
| Unknown_FAHFA 44:11 | 54947.72 | 82736.43 | 68270.17 | 130503.28 |
| Unknown_FAHFA 44:8 | 11454.44 | 9286.26 | 12068.88 | 21159.42 |
| Unknown_FAHFA 44:9 | 18069.24 | 23351.96 | 20003.92 | 26086.82 |
