## Supplemental Table S3 for "The MRL/MpJ Mouse Strain Is Not Protected From Muscle Atrophy And Weakness After Rotator Cuff Tear"

***Supplemental Table S3. qPCR Gene Expression.*** Gene expression values from infraspinatus muscles as measured with qPCR. Target gene expression is normalized to housekeeping gene *Csnk2a1* using the $2^{-\Delta Ct}$ approach. Values are mean±SD. Differences between groups assessed using a two-way ANOVA. Posthoc sorting: a, different (p<0.05) from B6 control; b, different (p<0.05) from B6 tear; c, different (p<0.05) from MRL control. N≥4 mice per group. Primer sequences are as follows (listed 5' to 3'): *Col1a1* forward, GGGGCAAGACAGTCATCGAA; *Col1a1* reverse, GGGTGGAGGGAGTTTACACG; *Csnk2a1* forward, GCCGCCATATTGTCTGTGTG; *Csnk2a1* reverse, CCCTTTTTCTTCACACTGCGG; *Fbxo32* forward, GCCCTCCACACTAGTTGACC; *Fbxo32* reverse, GACGGATTGACAGCCAGGAA; *Plin1* forward, GAGACTGAGGTGGCGGTCT; *Plin1* reverse, AACACCGAGAAAGCAGCTAGG; *Rpl14* forward, CGGGTGGCCTACATTTCCTT; and *Rpl14* reverse, GGGTCCATCCACTAAAGCCC.

| | **B6** | | **MRL** | |
|---|---|---|---|---|
| | *Control* | *Tear* | *Control* | *Tear* |
| *Col1a1* | 1.331±0.551 | 25.494±19.166[a] | 1.525±0.955[b] | 20.285±11.018[a,c] |
| *Fbxo32* | 0.975±0.470 | 0.079±0.042[a] | 0.706±0.145[b] | 0.043±0.015[a,c] |
| *Plin1* | 0.008±0.004 | 0.021±0.010 | 0.003±0.002 | 0.032±0.030[c] |
| *Rpl14* | 2.411±0.585 | 3.964±0.188[a] | 0.749±0.401[a,b] | 1.662±0.951[b,c] |
